# Cell type prioritization in single-cell data

**DOI:** 10.1101/2019.12.20.884916

**Authors:** Michael A. Skinnider, Jordan W. Squair, Claudia Kathe, Mark A. Anderson, Matthieu Gautier, Kaya J.E. Matson, Marco Milano, Thomas H. Hutson, Quentin Barraud, Aaron A. Phillips, Leonard J. Foster, Gioele La Manno, Ariel J. Levine, Grégoire Courtine

## Abstract

We present a machine-learning method to prioritize the cell types most responsive to biological perturbations within high-dimensional single-cell data. We validate our method, Augur (https://github.com/neurorestore/Augur), on a compendium of single-cell RNA-seq, chromatin accessibility, and imaging transcriptomics datasets. We apply Augur to expose the neural circuits that enable walking after paralysis in response to spinal cord neurostimulation.

---

Within a decade, single-cell technologies have scaled from individual cells to entire organisms^1,2^. Investigators are now able to quantify RNA and protein expression, resolve their spatial organization in complex tissues, and dissect their regulation in hundreds of thousands of cells. This exponential increase in scale is enabling a transition from atlasing of healthy tissues to delineating the cell type-specific responses to disease and experimental perturbation^3–5^. This shift requires a parallel analytical transition, from cataloging the marked molecular differences between cell types to resolving more subtle phenotypic alterations within cell types. Existing tools focus on identifying individual genes or proteins with statistically significant differences between conditions^6^. However, inferences at the level of individual analytes are ill-suited to address the broader question of which *cell types* are most responsive to a perturbation in the multidimensional space of single-cell data. We propose that such prioritizations could clarify the contribution of each cell type to organismal phenotypes such as disease state, or identify cellular subpopulations that mediate the response to external stimuli such as drug treatment. Cell type prioritization could also guide downstream investigation, including the selection of experimental systems such as Cre lines or FACS gates to support causal experiments.

Here, we introduce Augur, a versatile method to prioritize cell types based on their molecular response to a biological perturbation (**Fig. 1a**). We reasoned that cell types most responsive to a perturbation should be more separable than less affected ones. In turn, we hypothesized that the relative difficulty of this separation would provide a quantitative basis for cell type prioritization. We formalized this difficulty as a classification task, asking how accurately disease or perturbation state could be predicted from highly multidimensional single-cell measurements. For each cell type, Augur withholds a proportion of sample labels, and trains a classifier on the labeled subset. The classifier predictions are compared with the experimental labels, and cell types are prioritized based on the area under the receiver operating characteristic curve (AUC) of these predictions in cross-validation.

**Fig. 1.**
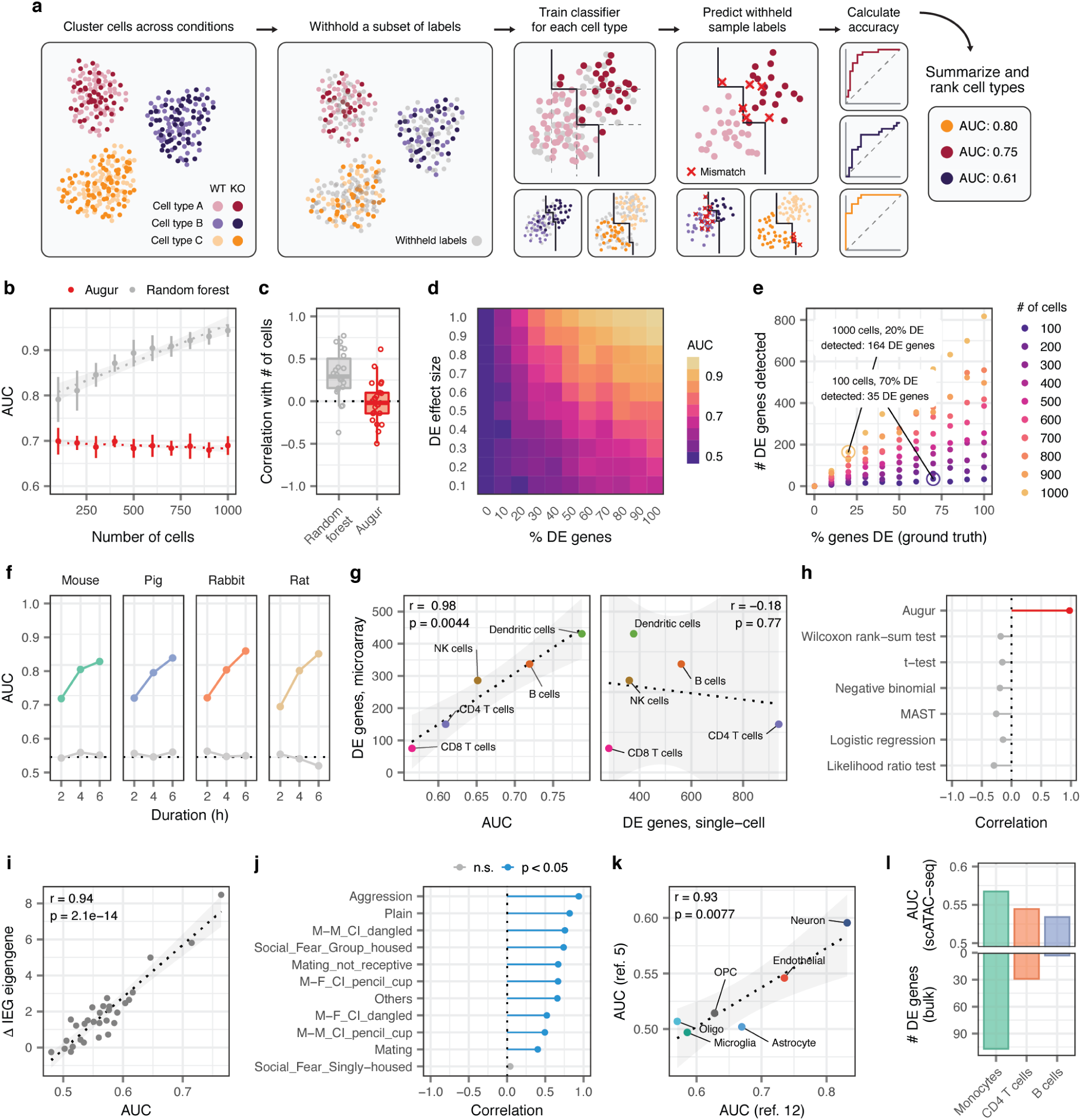
Augur correctly prioritizes cell types in synthetic and experimental single-cell datasets. **a**, Schematic overview of Augur. **b**, AUCs of Augur and a naive random forest classifier without subsampling in simulated scRNA-seq datasets containing increasing numbers of cells. Cell type prioritizations are confounded by training dataset size for the naive classifier, but Augur abolishes this confounding factor. The mean and standard deviation of ten simulation replicates are shown. **c**, Pearson correlations between the AUC of each cell type, and the number of cells of that type sequenced, across a compendium of 22 scRNA-seq datasets, for Augur and a naive random forest classifier without subsampling. **d**, Augur AUCs scale monotonically with both the proportion of DE genes and the magnitude of DE in simulated cell populations. **e**, Relationship between number of DE genes detected by a representative test for single-cell differential gene expression (Wilcoxon rank-sum test), and the proportion of differentially expressed genes simulated between the two populations, for simulated populations of between 100 and 1,000 cells. **f**, Augur cell type prioritizations track with duration of LPS exposure in a cross-species scRNA-seq experiment^7^. Grey points show AUCs with sample labels randomly permuted. **g-h**, Cell type prioritization in matched single-cell^4^ and bulk^8^ transcriptomic profiles of PBMCs after interferon stimulation. **g**, Left, Augur cell type prioritizations mirror the number of DE genes in a microarray dataset of FACS-purified cells. Right, the number of DE genes detected in the scRNA-seq dataset by a Wilcoxon rank-sum test is uncorrelated with the FACS gold standard. **h**, Correlation coefficients between cell type prioritizations (AUC or number of DE genes) in the scRNA-seq dataset and the FACS gold standard. **i-j**, Cell type prioritization in the mouse ventromedial hypothalamus reflects induction of IEG transcription. **i**, Correlation between AUC and the difference in the first principal component of IEG expression (ΔIEG eigengene) engaging in aggressive behavior. **j**, Pearson correlation coefficients between cell type-specific AUC and ΔIEG eigengene values for eleven behavioral stimuli^9^. **k**, Reproducibility of cell type prioritization in two independent scRNA-seq studies of Alzheimer’s disease^5,10^. **l**, Augur cell type prioritizations in a scATAC-seq dataset^11^ track with the number of DE genes in an RNA-seq dataset of FACS-purified cells.

Because the amount of available training data typically has a strong effect on classifier performance, we anticipated that the uneven relative abundances of cell types in single-cell datasets could confound cell type prioritization. In both simulated data and a compendium of 22 published scRNA-seq datasets, we found that the AUC scaled with the number of cells, as opposed to the perturbation intensity (**Supplementary Figs. 1a-b** and **2**). To overcome this confounding factor, Augur repeatedly draws small samples from the dataset, and reports the mean AUC across samples. We found this procedure abolished the dependence on the total number of cells (**Fig. 1b-c, Supplementary Fig. 1c-d**, and **Supplementary Fig. 2**). Moreover, we established that Augur correctly prioritized cell types subjected to simulated perturbations of known intensities, finding the AUC increased monotonically with both the amount and magnitude of simulated differential expression (**Fig. 1d** and **Supplementary Fig. 1e-f**).

Prior studies have attempted to prioritize cell types based on the relative number of genes passing a statistical threshold for differential expression (DE)^5,12^. In both simulated and experimental datasets, however, we found the number of DE genes was strongly correlated with the number of cells per type (**Supplementary Figs. 1g-i** and **2c**), causing abundant cell types with modest transcriptional perturbations to be prioritized over rare but more strongly perturbed cell types (**Fig. 1e** and **Supplementary Fig. 1j**).

We applied Augur to several scRNA-seq datasets in order to evaluate its ability to prioritize cell types involved in well-understood biological processes. Augur detected the expected dose-response relationship in bone marrow-derived mononuclear phagocytes from four species stimulated with LPS for between two and six hours, with similar AUCs across species^7^ (**Fig. 1f**). We observed similar results for Jurkat cells stimulated with PMA/ionomycin^13^ (**Supplementary Fig. 3a**). We next applied Augur to a scRNA-seq dataset of PBMCs stimulated with interferon^4^, comparing cell type prioritizations to an independent microarray experiment on FACS-purified cells^8^. We observed an almost perfect correspondence between Augur and the number of DE genes in this FACS gold standard (**Fig. 1g**). In contrast, the number of differentially expressed genes in the scRNA-seq dataset was weakly or negatively correlated with the gold standard (**Fig. 1g-h** and **Supplementary Fig. 3b**). Finally, we applied Augur to prioritize neuron subtypes of the ventromedial hypothalamus in response to various behavioral stimuli^9^. Augur prioritizations were correlated with the relative induction of intermediate early gene (IEG) transcription across a range of social behaviors (**Fig. 1i-j** and **Supplementary Fig. 3c**).

We then evaluated the reproducibility of cell type prioritization by applying Augur to two independent scRNA-seq studies comparing individuals with Alzheimer’s disease and healthy controls^5,10^. Augur produced nearly identical prioritizations, identifying the most profound transcriptional perturbations in neurons and endothelial cells (**Fig. 1k**). Similarly, we asked whether Augur could prioritize cell types from identical experimental perturbations, but obtained with orthogonal single-cell technologies. We applied Augur to scRNA-seq^14^ and single-cell imaging transcriptomics (STARmap)^15^ datasets from the visual cortex after exposure to light. Despite technical differences between the datasets, Augur consistently prioritized excitatory neurons, and even ranked subpopulations of excitatory neurons from specific cortical layers in identical order (**Supplementary Fig. 3d**). Finally, we applied Augur to single-cell ATAC-seq data from bone marrow-derived cells stimulated with LPS^11^, and found that Augur cell type prioritizations mirrored a gold standard from bulk RNA-seq of FACS-sorted cells (**Fig. 1l**)^16^.

Augur can flexibly incorporate continuous or multi-class sample labels in addition to conventional treatment versus control designs. We applied Augur to prioritize cell types of the prefrontal cortex based on quantitative measures of amyloid burden, neuritic plaques, and neurofibrillary tangles in individuals with Alzheimer’s disease^5^. Cell type prioritizations were strongly correlated to those based on clinical diagnosis, reflecting the pathogenesis of the disease (**Supplementary Fig. 4**). Likewise, Augur can readily be applied to prioritize cell types in datasets with more than two perturbations (**Supplementary Fig. 5**).

To apply Augur to single-cell datasets with more complex experimental designs, we devised a test for differential cell type prioritization (**Supplementary Fig. 6a**). Applying differential prioritization to a single-cell imaging transcriptomics (MERFISH) dataset^17^, Augur identified multiple neuron subtypes preferentially activated during parenting in either male or female mice (**Supplementary Fig. 6b-c**). Similarly, in a scRNA-seq dataset^18^, Augur prioritized several neuron subtypes with differential responses to whisker lesioning in Cx3cr1^+/−^ and Cx3cr1^−/−^ mice (**Supplementary Fig. 6d**).

We also considered whether Augur could be applied directly to single-cell measures of transcriptome dynamics, such as the RNA velocity^19^, in order to specifically prioritize cell types undergoing an acute response to a perturbation on the timescale of transcription. We found that both experimental measurements^20^ and computational inference^19^ of transcriptional activity consistently captured more information than total RNA abundance in perturbations ranging from 45 min to 4 h in duration (**Supplementary Fig. 7a-g**). Conversely, we confirmed that transcriptome dynamics did not confer an appreciable information gain to cell type prioritization when the perturbation is chronic (**Supplementary Fig. 7h-i**).

We finally aimed to demonstrate the relevance of Augur to discover new biological mechanisms. We^21^ and others^22,23^ have shown that targeted epidural electrical stimulation of the lumbar spinal cord (TESS), augmented by monoaminergic stimulation^24^, restores walking after spinal cord injury in individuals with paralysis. However, the neural circuits engaged by this treatment remain enigmatic. We devised an experiment to expose the neuron subtypes recruited by TESS using single-cell transcriptomics (**Fig. 2a**). Mice received a severe contusion of the thoracic spinal cord that led to permanent paralysis of both legs. In the presence of serotonergic and D1 agonists, TESS immediately enabled walking in paralyzed mice (**Fig. 2b-c**). We performed single-nucleus RNA-seq of 18,514 nuclei from mice walking for 30 min with TESS and control mice, identifying all the major cell types of the lumbar spinal cord (**Fig. 2d** and **Supplementary Fig. 8**). We then subjected the 6,035 identified neurons to an additional round of clustering. This analysis identified 39 neuron subtypes expressing classical marker genes and were detected across experimental conditions (**Fig. 2e** and **Supplementary Fig. 9**).

**Fig. 2.**
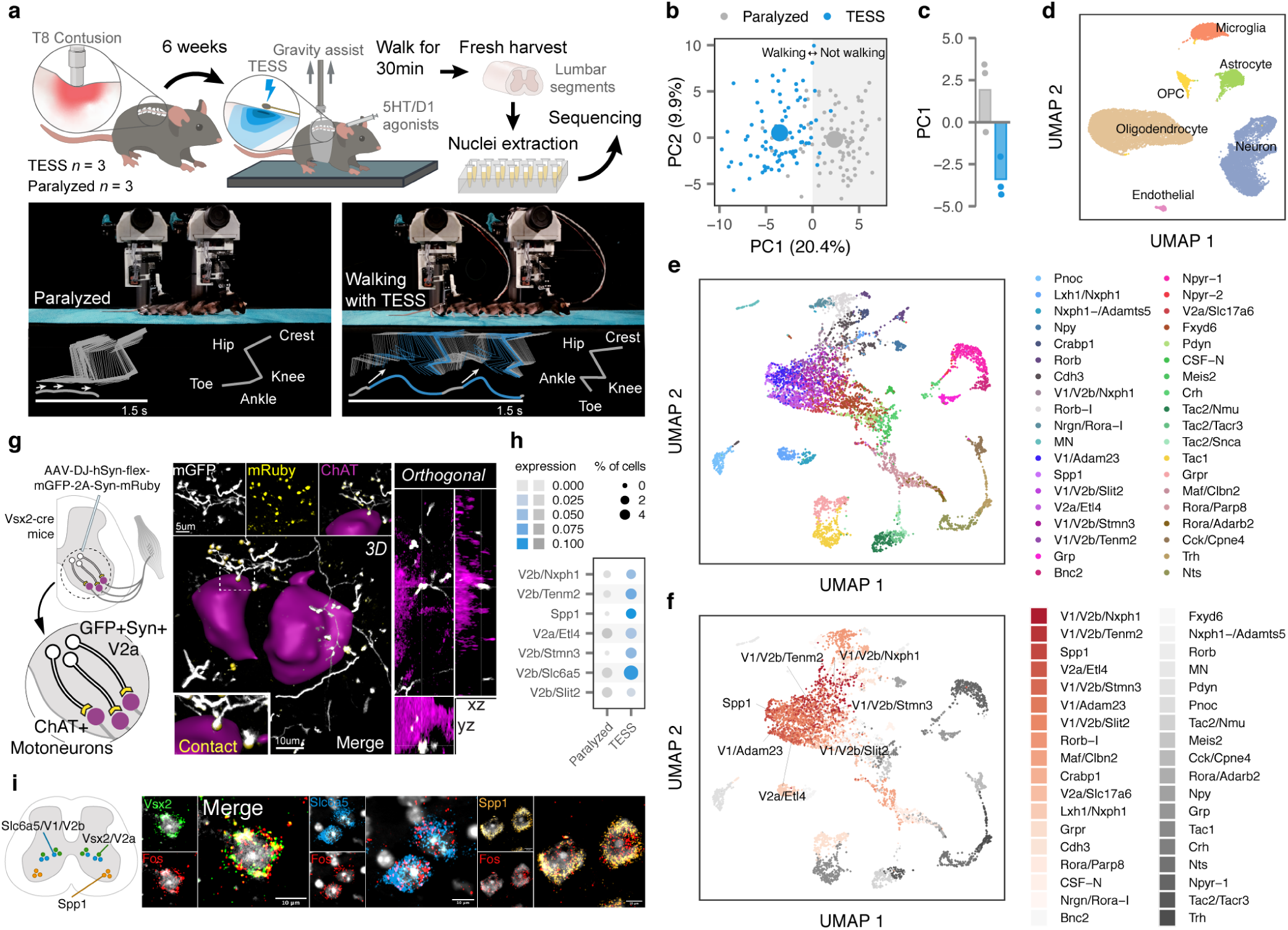
Augur identifies neuron subtypes that enable walking after paralysis. **a**, Top, single-nucleus RNA-sequencing experimental design to prioritize neuron subtypes recruited by TESS. Middle, chronophotography of mice in the presence or absence of TESS and monoaminergic agonists. Bottom, stick diagram decompositions of right leg movements; leg endpoint trajectory with acceleration at toe-off; activity of extensor and flexor muscles of the ankle. **b**, Principal component analysis of gait parameters for each condition (small circles). Large circles show the average per group. **c**, Bar plot shows the average scores on principal component 1 (PC1), which quantify the locomotor performance of paralyzed mice (*n* = 3) and mice walking with TESS (*n* = 3). **d**, Uniform manifold approximation and projection (UMAP) visualization of 18,514 nuclei, revealing the six major cell types of the mouse lumbar spinal cord. **e**, UMAP visualization of 6,035 neurons subjected to an additional round of sub-clustering and the 39 identified neuron subtypes. **f**, UMAP neuron visualization, colored by Augur cell type prioritization (AUC). The seven prioritized neuron subtypes with the highest AUCs are highlighted. **g**, Monosynaptically restricted anterograde tracing in Vsx2-Cre mice reveals V2a interneurons densely innervating motor neurons (ChAT). **h**, Dot plot showing expression of the immediate early gene Fos in neuron subtypes prioritized by Augur. **i**, Confirmation of colocalization of V2a, V1/V2b, and Spp1 marker genes (Vsx2, Slc6a5, and Spp1 respectively) and Fos by RNAscope *in situ* hybridization. Schematic indicates location of imaging for each marker within the spinal cord.

We reasoned that applying Augur directly to the RNA velocity of these neurons could prioritize subtypes that are immediately engaged by the therapy. Previous studies suggested that TESS generates an electrical field that depolarises proprioceptive afferent fibers^25^. Consistent with this prediction, Augur robustly prioritized interneurons with the molecular profiles of V2a and V1/V2b neurons, which are known to receive synapses from proprioceptive afferents (**Fig. 2f** and **Supplementary Fig. 10**). V2a interneurons have been implicated in left-right alternation^26^, whereas V2b interneurons are critical for flexor-extensor alternation^27^. Augur also prioritized Spp1-positive neurons, typically associated withs motoneurons (**Fig. 2f**). Virus-mediated anatomical tracing in transgenic mice revealed dense synaptic projections from the prioritized interneurons onto motoneurons (**Fig. 2g**). The pronounced induction of immediate early genes in V2a and V1/V2b interneurons (**Fig. 2h**) confirmed their activation in response to TESS-enabled walking, a finding we verified by *in situ* hybridization (**Fig. 2i** and **Supplementary Fig. 11**). In contrast, interneurons not prioritized by Augur showed minimal amounts of Fos mRNA (**Supplementary Fig. 11**). These results illustrate the value of Augur to expose neural circuits underlying complex behaviors.

Augur is computationally efficient, requiring a median of 49.7 min and 2.3 GB of RAM to analyze our compendium of 22 scRNA-seq datasets (**Supplementary Fig. 12a-b**). Inherent to the design of Augur is the ability to scale to datasets containing hundreds of thousands or even millions of cells on a laptop (**Supplementary Fig. 12c-d**). Moreover, Augur is robust to sequencing depth and classifier hyperparameters (**Supplementary Figs. 13-14**). As an efficient and principled method for cell type prioritization, we envision that Augur will facilitate the interpretation of a growing resource of single-cell data spanning multiple experimental conditions, and help single-cell technologies realize their potential to pinpoint cell types underlying organism-level phenotypes.

## Methods

### Design and implementation of Augur

Single-cell technologies increasingly allow investigators to collect datasets that span multiple experimental conditions: for instance, patients with a particular disease compared to healthy controls, animals exposed to a specific behavioral stimulus compared to unstimulated animals, or organisms subject to a particular genetic manipulation and wild-type controls. A number of tools have been developed to identify individual analytes (for instance, genes, proteins, or accessible chromatin regions) that exhibit statistically significant differences between experimental conditions^6,28^. However, for many biological questions, the analytical level of interest is not individual differentially abundant features, but rather the specific cell types that are most strongly affected by a stimulus. For instance, investigators may design a single-cell transcriptomics experiment to identify particular cell types in a complex tissue that undergo the most marked transcriptional changes in response to treatment with a drug, in order to clarify its mechanism of action. We refer to the process of ranking cell types based on their molecular response to a biological perturbation as cell type prioritization.

We designed Augur as a method to prioritize cell types based on their molecular response to a perturbation in highly multidimensional single-cell data. We reasoned that cells undergoing a profound response to a given experimental stimulus should become more separable, in the space of molecular measurements, than cells that remain unaffected by the stimulus. We sought to design a quantitative metric of this separability that would be robust to heteroscedasticity between cell types, and account for the specific biological and technical variability within each cellular subpopulation. Accordingly, Augur quantifies this separability by asking how readily the experimental sample labels associated with each cell (e.g., treatment vs. control) can be predicted from molecular measurements alone. In practice, this is achieved by training a machine-learning model specific to each cell type, to predict the experimental condition from which each individual cell originated. The accuracy of each cell type-specific classifier is evaluated in cross-validation, providing a quantitative basis for cell type prioritization.

We reasoned that an ideal method for cell type prioritization would make no assumptions about the distributions of features provided as input^29^, and more broadly, would be agnostic to the particular molecular features provided as input: that is, it would readily incorporate single-cell RNA-seq^30–33^, proteomics^34,35^, epigenomics^11,36–38^, and imaging transcriptomics^15,17,39^ datasets, among other modalities. Accordingly, Augur uses a random forest^40^ classifier to predict sample labels for each cell type. Random forests have the advantage that they do not make any parametric assumptions about the distribution of the input features, and consequently are robust to both the nature of the molecular measurements themselves, as well as to the specific pre-processing and normalization steps applied to obtain the input features-by-cells matrix.

When training machine-learning models, model performance generally improves as the size of the training dataset increases. We anticipated that this well-known phenomenon could present a critical confound to cell type prioritization, because cell types are unevenly represented in most single-cell datasets for both biological and technical reasons. To account for this confound, Augur repeatedly draws small samples of fixed size from each cell-type specific gene expression matrix, and performs cross-validation on these sub-sampled matrices (by default, 50 subsamples of 20 cells per condition are drawn). Augur then reports the mean cross-validation AUC across many small subsamples. We confirmed that this procedure abolishes the relationship between the number of cells of a particular type and the cross-validation AUC, in both real and simulated datasets (**Fig. 1b-c and Supplementary Figs. 1-2)**.

To further improve computational efficiency, Augur incorporates two feature selection steps to minimize the number of analytes provided to the classifier as input. First, for each cell type in turn, Augur removes genes with little cell-to-cell variation within that cell type. This procedure, commonly referred to as highly variable gene identification in the context of single-cell RNA-seq^41^, also has the effect of removing noise. To flexibly account for the mean-variance relationship without making assumptions about the form of this relationship, Augur fits a local polynomial regression between the mean and coefficient of variation^42,43^ using the ‘loess’ function, and ranks genes based on their residuals in this model. A fixed quantile of the most highly variable genes are retained for each cell type (specified using the ‘var_quantile’ parameter, which defaults to 50% in order to remove only features that show less-than-expected variation based on their mean abundance). Second, for each iteration, a random proportion of features are randomly removed to improve speed and memory usage (specified using the ‘feature_perc’ parameter, which also defaults to 50%). In combination, these steps significantly reduce the size of the matrix that must be taken out of a sparse representation for input to the classifier, from ∼20,000 genes to ∼5,000 genes in a typical scRNA-seq dataset. To avoid discarding information in datasets where fewer analytes are measured, feature selection is only performed for datasets exceeding a certain minimum number of features (with this cutoff set, by default, to 1,000).

#### Implementation

Augur is implemented as an R package, available from https://github.com/neurorestore/Augur (**Supplementary Fig. 15)**. Augur takes as input a features-by-cells (e.g., genes-by-cells for scRNA-seq) matrix, and a data frame containing metadata associated with each cell, minimally including the cell type annotations and sample labels to be predicted. Alternatively, a Seurat^44^, monocle3^45^, or SingleCellExperiment^46^ object can be provided as input. To optimize both speed and memory usage, all computations are implemented for sparse matrices, up to the classification procedure itself. Because the feature selection, classification, and cross-validation procedures are independent for each cell type, Augur can readily be parallelized over the cell types in the input dataset, using the ‘mclapply’ package for parallelization, and runs on four cores by default.

#### Multiclass classification and regression

Augur quantifies the accuracy by which cell type labels can be predicted from molecular measurements using the area under the receiver operating characteristic curve (AUC), or the macro-averaged AUC in the case of multiclass classification. For experiments in which the perturbation can be interpreted as a continuous or ordinal variable, the classification objective is replaced with a regression task, and the accuracy of the corresponding random forest regression models is quantified using the concordance correlation coefficient (CCC)^47^, a measure of both the precision and accuracy of the relationship between predicted and experimental sample labels. By default, Augur returns the mean AUC (or CCC) for each cell type as a summary of cell type classification, but also calculates a larger suite of metrics for each fold of each subsampling iteration, including accuracy, precision, recall, sensitivity, specificity, negative predictive value, and positive predictive value, for users interested in investigating predictions in more detail.

#### Differential prioritization

To compare cell type prioritizations between related conditions, we devised a permutation-based test for differential prioritization. In order to obtain a null distribution of AUCs for each cell type that reflected variability associated with number of cells sequenced, read depth, and other technical factors, we permuted sample labels within each cell type, and ran Augur on the permuted dataset. We repeated this permutation procedure 1,000 times. We then compared the observed difference between condition-specific AUCs, ΔAUC_obs_, for each cell type to the difference under permuted sample labels, ΔAUC_rnd_, and calculated permutation p-values^48^.

### Simulations

We initially tested Augur on simulated scRNA-seq data, using the ‘Splatter’ R package^49^. Initial simulation parameters were estimated from the Kang et al. dataset^4^ using the ‘splatEstimate’ function, and populations of 100–1,000 cells from two experimental conditions were generated, in increments of 100. We then simulated differential expression in varying proportions of genes (using the ‘de.prob’ parameter), and with varying magnitudes (using the ‘de.facLoc’ parameter). To specifically evaluate the ability of Augur to abolish the relationship between the number of cells in a particular population and the AUC of sample label classification, we compared Augur to cell type prioritization using an identical feature selection and classification procedure, but without drawing small subsamples from the dataset, by setting the ‘n_subsamples’ argument to 0. We additionally implemented a cell type prioritization scheme based on the number of differentially expressed genes between conditions, as previously described^5,12^. Cell types were ranked based on the number of differentially expressed genes using six different tests for differential expression in single-cell transcriptomics datasets (t-test, Wilcoxon rank-sum test, likelihood ratio test^50^, logistic regression^51^, MAST^52^, and a negative binomial generalized linear model), implemented through the Seurat ‘FindMarkers’ function.

### RNA velocity analysis

To generate intronic and exonic read count matrices for each dataset, data were downloaded from the SRA and converted to FASTQ format using the SRA toolkit. In the case of inDrops^14^ data, annotated BAM files were obtained using dropTag^53^ with flags -s -S -c. Reads were then aligned to the latest Ensembl release (GRCm38.93), using STAR (v.2.5.3a)^54^. For Drop-seq data^12,55^, files were first converted from FASTQ to BAM format using the Picard function ‘FastqtoSam’. Reads were then aligned to the latest Ensembl release using the Drop-seq toolkit (https://github.com/broadinstitute/Drop-seq)^56^. Next, count matrices of exonic and intronic reads were obtained using dropEst^53^ with flags -m -V -L eEBA -F. Barcodes were filtered to match those present in the processed datasets uploaded to the Gene Expression Omnibus (GEO) for each dataset. RNA velocity was subsequently calculated using the ‘velocyto’ R package^19^. Features were first chosen by filtering for genes with a minimum expression value per cell type using the function ‘filter.genes.by.cluster.expression’, with filters adjusted based on the read count distributions for each dataset (GSE102827: exon filter, 0.5, intron filter, 0.1; GSE130597: exon filter, 0.03, intron filter, 0.02; GSE103976, exon filter, 0.05, intron filter, 0.03). We then calculated gene-relative velocity using kNN pooling with k = 10 (default) and fit.quantile = 0.01. By default, the function ‘gene.relative.velocity.estimates’ in velocyto.R returns a matrix containing only those features for which accurate estimates of *γ* and velocity could be obtained. Consequently, we ran Augur without either variable gene or random gene filters, as feature selection had already been performed during the creation of the RNA velocity matrix used as input. To compare AUCs for cell type prioritization on matrices of exonic or total counts, we retained only those genes for which velocity estimates could be calculated, and likewise disabled the variable gene and random gene filters. All other parameters were left as default.

### Computational benchmarking

To quantify the computational resources required for cell type prioritization (**Supplementary Fig. 12**), we ran Augur with default settings on our compendium of 22 scRNA-seq datasets. The R package ‘peakRAM’ was used to monitor peak memory usage, and the base R function ‘system.time’ was used to monitor wall time.

### Hyperparameter analysis

To characterize the robustness of Augur prioritizations to hyperparameters associated with its sub-sampling or feature selection procedures, the random forest classifier, and the choice of classifier itself, we evaluated the impact of systematically varying each of these parameters (**Supplementary Fig. 13**). We first investigated the impact of the number and size of subsamples from each cell-type-specific gene expression matrix on cell type prioritization, finding the ranks of each cell type stabilized around 50 subsamples. While larger subsample sizes generally yielded more robust ranks, these thresholds also precluded analysis of several cell types represented by fewer cells in existing datasets, and consequently we opted for an inclusive subsample size of 20 cells per experimental condition. Similarly, we ran Augur on gene expression matrices consisting of the top 10–100% of highly variable genes, followed by selection of a random subset of 10–100% of these, but found Augur was generally robust to the features provided as input. (We used the default thresholds of 50% on the variable gene and random selection filters throughout, unless otherwise specified). To assess the robustness of Augur prioritizations to random forest hyperparameters, we varied the number of trees in the forest between 10–1,000, the minimum number of cells required to split an internal node between 2–10, and the number of features sampled per split between 2–500. Finally, to assess the impact of the classifier itself, we implemented L1-penalized logistic regression in Augur using the R package ‘glmnet’, with the optimal value of the regularization parameter *λ* determined for each iteration using the function ‘cv.glmnet’.

### Downsampling analysis

Motivated by the observation that only a fraction of reads at conventional depths are required to detect transcriptional programs and assign cell types^57^, we also evaluated the impact of sequencing depth on Augur cell type prioritizations by downsampling published scRNA-seq datasets to between 5–95% of their original depths (**Supplementary Fig. 14**). Reads were down-sampled from the processed count matrices using the ‘downsampleMatrix’ function from the ‘DropletUtils’ package^58^.

### Preprocessing and analysis of published single-cell datasets

Data from a total of 29 published single-cell studies was processed and analyzed with Augur as described below. Unless otherwise noted, expression matrices and metadata were stored as Seurat objects, and genes detected in less than three cells were removed.

*Arneson et al., 2018*^59^. scRNA-seq data from the hippocampus of mice after a mild traumatic brain injury (mTBI), delivered using a mild fluid percussion injury model, and matched controls was obtained from GEO (accession: GSE101901). Metadata, including cell type annotations, was provided by the authors. AUCs were calculated by comparing cells from mTBI and control mice.

*Avey et al., 2018*^60^. scRNA-seq data from the nucleus accumbens of mice treated with morphine for 4 h and saline-treated controls was obtained from GEO (accession: GSE118918). Cells identified as doublets and non-unique barcodes were removed. Metadata, including cell type annotations, was provided by the authors. AUCs were calculated by comparing cells from morphine- and saline-treated mice.

*Aztekin et al., 2019*^61^. scRNA-seq data from regeneration-competent (NF stage 40-41) Xenopus laevis tadpoles was obtained from ArrayExpress (E-MTAB-7716). AUCs were calculated by comparing cells from tadpoles at 1 d post-amputation to control tadpoles.

*Cheng et al., 2019*^62^. scRNA-seq data from intestinal crypt cells in wild-type and Hmgcs2 knockout mice was obtained directly from the authors of the original publication. AUCs were calculated by comparing wild type and control mice.

*Erhard et al., 2019*^20^. scSLAM-seq data from CMV-infected or uninfected fibroblasts was obtained from GEO (accession: GSE115612), including both total count and new-to-total RNA ratio (NTR) matrices as calculated by the authors. In cases where the NTR could not be estimated, missing values were imputed as the median value. AUCs were calculated by comparing cells from CMV-infected and uninfected fibroblasts.

*Grubman et al., 2019*^10^. scRNA-seq data from post-mortem entorhinal cortex of patients with Alzheimer’s disease and matched controls was obtained directly from the authors prior to publication. Cells annotated as ‘undetermined’ and ‘doublet’ were removed prior to downstream analysis. AUCs were calculated by comparing cells from individuals with Alzheimer’s disease and control individuals.

*Gunner et al., 2019*^18^. scRNA-seq data from the barrel cortex before or after whisker lesioning (sensory deprivation) in Cx3cr1^+/−^ and Cx3cr1^−/−^ mice was obtained from GEO (accession: GSE129150). Cell types not included in **Supplementary Fig. 10** of the original publication were removed. AUCs were calculated by comparing cells from deprived and control animals, in Cx3cr1^+/−^ and Cx3cr1^−/−^ mice separately. Differential prioritization was applied to identify neuron subtypes preferentially affected by whisker lesioning in mice of either genotype. For the analysis of confounding factors presented in **Supplementary Fig. 2**, only cells from homozygous mice were used.

*Haber et al., 2017*^3^. scRNA-seq data from epithelial cells of the mouse small intestine in healthy mice and after two days of Salmonella infection was obtained from GEO (accession: GSE92332), using the Drop-seq data collected by the original publication. AUCs were calculated by comparing cells from infected and uninfected mice.

*Hagai et al., 2018*^7^. scRNA-seq data from bone marrow-derived mononuclear phagocytes from four different species (rat, rabbit, pig, mouse) exposed to lipopolysaccharide (LPS) for 2, 4, or 6 h was obtained from ArrayExpress (accession: E-MTAB-6754). Cells whose total number of counts was above the 97.5th percentile were excluded as possible doublets. AUCs were calculated by comparing cells from each LPS-stimulated timepoint (2, 4, and 6 h) to a common population of unstimulated controls in each species separately.

*Hrvatin et al., 2018*^14^. scRNA-seq data from the visual cortex of mice housed in darkness, then exposed to light for 0 h, 1 h, or 4 h was obtained from GEO (accession: GSE102827). Cell types labeled as ‘NA’ were removed from downstream analyses, as were three small populations of cells from the subiculum, hippocampus, and retrosplenial cortex, for consistency with the original publication. AUCs were calculated at each timepoint separately by comparing cells in each light exposure group (1 h or 4 h) to the common population of unexposed control cells. Cell types were compared to those defined by the three-dimensional intact-tissue sequencing method STARmap^15^ by averaging gene expression profiles over the 139 genes quantified in both experiments, then taking the Spearman correlation between average gene expression profiles for each cell type. We also calculated macro-averaged AUCs in a multiclass classification task incorporating cells stimulated with light for 0 h, 1 h, or 4 h, and compared AUCs and permutation p-values between multiclass classification and each of the three possible pairwise binary classification tasks (i.e., 0 h vs. 1 h, 0 h vs. 4 h, and 1 h vs. 4 h). The 4 h comparison was used for the analysis of confounding factors presented in **Supplementary Fig. 2**. For RNA velocity analysis, raw sequencing data was obtained from the SRA (accession: PRJNA399082) and processed as described above.

*Hu et al., 2017*^63^. snRNA-seq from the cerebral cortex of mice after pentylenetetrazole (PTZ)-induced seizure and saline-treated controls was obtained from the Google Drive folder accompanying the original publication (https://github.com/wulabupenn/Hu_MolCell_2017). AUCs were calculated by comparing cells from PTZ- and saline-treated mice.

*Jaitin et al., 2019*^64^. scRNA-seq data from white adipose tissue of mice fed either a high-fat diet or normal chow for six weeks were obtained from the Bitbucket repository accompanying the original publication (https://bitbucket.org/account/user/amitlab/projects/ATIC). Metadata, including cell type annotations, were provided by the authors. AUCs were calculated by comparing cells from high-fat diet and normal chow-fed mice.

*Kang et al., 2018*^4^. scRNA-seq data from peripheral blood mononuclear cells (PBMCs) stimulated with recombinant *IFN-β* for 6 h and unstimulated PBMCs was obtained from GEO (accession: GSE96583). Doublets and unclassified cells were removed. AUCs were calculated for each of the eight distinct cell types (CD14+ monocytes, CD4 T cells, dendritic cells, NK cells, CD8 T cells, B cells, megakaryocytes, and FCGR3A+ monocytes) by comparing IFN-stimulated and unstimulated cells. These AUCs were subsequently compared to a bulk (microarray) dataset obtained from FACS-sorted mouse PBMC populations exposed to purified IFN-*α*, 2 h after subcutaneous injection^8^. Differentially expressed genes were obtained from the original publication. For this comparison, only the five cell types included in both studies were examined. We additionally compared the number of genes called as differentially expressed in each cell type within the scRNA-seq data to the bulk gold standard using six different tests for differential expression, as described above.

*Kim et al., 2019*^9^. scRNA-seq data from the ventromedial hypothalamus of mice exposed to one of eleven behavioral stimuli and control mice was obtained from the Mendeley repository accompanying the original publication. Cell type annotations were provided directly by the authors. For each behaviour, AUCs were calculated by comparing cells from animals engaging in the behaviour to the common population of control animals. IEG ‘eigengenes’ were calculated as the first principle component of IEG expression^65^, using a previously published list of immediate early genes^14^, and the difference in average IEG eigengene between conditions per cell type was compared to the AUC. For the analysis of confounding factors presented in **Supplementary Fig. 2**, only the aggression condition was used.

*Lareau et al., 2019*^11^. scATAC-seq data from resting and LPS-stimulated bone marrow cells was obtained from GEO (accession: GSE123580). AUCs for monocytes, CD4 T cells, and B cells were calculated by comparing stimulated and unstimulated cells, and compared to the number of genes called as differentially expressed in an independent bulk RNA-seq experiment using a low-input microfluidic platform to analyze FACS-purified cells^16^. Differentially expressed genes were obtained from the original publication.

*Mathys et al., 2019*^5^. snRNA-seq data from post-mortem pre-frontal cortex of patients with Alzheimer’s disease and matched controls was obtained from Synapse (accession: syn18681734). Patient data and additional metadata were also obtained from Synapse (accessions: syn3191087 and syn18642926, respectively). For the comparison to the Grubman et al. 2019 dataset^10^, the coarse-grained clustering level was used, annotated as ‘broad.cell.type’. Excitatory and inhibitory neurons were combined into a single category of neurons in order to match the labels between datasets. AUCs were then calculated for each coarse-grained cell type by comparing cells from Alzheimer’s disease and control patients. For regression analyses, CCCs were calculated using three continuous, neuropathologically determined patient outcomes as sample labels, including neurofibrillary tangles (‘nft’), neuritic plaques (‘plaq_n’), and amyloid burden (‘amyloid’).

*McGinnis et al., 2019*^13^. scRNA-seq data from Jurkat T cells treated with PMA/ionomycin for between 15 min and 24 h was obtained from GEO (accession: GSE129578), with metadata inferred from MULTI-seq barcodes obtained (including cell types and stimulus duration for each cell) obtained from the supplementary materials accompanying the original publication. Mouse cells and genes and unstimulated human embryonic kidney cells profiled in the same experiment were removed. AUCs were calculated by comparing cells from each PMA/ionomycin-stimulated timepoint (15 min, 30 min, 1 h, 2 h, 4 h, 6 h, and 24 h) to a common population of unstimulated controls in each species separately.

*Moffitt et al., 2018*^17^. MERFISH data from the hypothalamic preoptic region in male and female mice displaying parenting behaviors and control mice was obtained from the Dryad repository accompanying the original publication. Cell types were assigned as specific neuron subtypes for neurons, and coarse cell type labels otherwise. Cells identified as putative doublets were removed. Five blank control barcodes were also removed, as was Fos, which was not measured in several animals. AUCs were calculated by comparing cells from parenting and control mice, after which differential prioritization was applied to identify neuron subtypes preferentially activated during parenting behaviors in male or female mice. Additionally, to assess whether the neuron subtypes that displayed sex-specific responses to parenting were also transcriptionally distinct in control animals, we used Augur to prioritize cell types in the naive population, using animal sex as the sample label.

*Ordovas-Montanes et al., 2018*^66^. scRNA-seq data from ethmoid sinus cells of patients with chronic rhinosinusitis (CRS), with and without nasal polyps, was obtained from **Supplementary Table 2** of the original publication. AUCs were calculated by comparing cells from patients with polyposis and non-polyposis CRS.

*Rossi et al., 2019*^12^. scRNA-seq data from the hypothalamus of mice fed either a high-fat diet or normal chow for between 9-16 weeks was obtained directly from the authors, in the form of a processed Seurat^44^ object. Cells annotated as ‘unclassified’ were removed. AUCs were calculated by comparing cells from high-fat diet and normal chow-fed mice. For RNA velocity analysis, raw sequencing data was obtained from the SRA (accession: PRJNA540713) and processed as described above.

*Sathyamurthy et al., 2018*^67^. snRNA-seq data from the spinal cord parenchyma of adult mice exposed to formalin or matched controls was obtained from GEO (accession: GSE103892). Cell types with blank annotations, or annotated as ‘discarded’, were removed. AUCs were calculated by comparing cells from mice exposed to formalin and control animals.

*Schirmer et al., 2019*^68^. snRNA-seq data from cortical and subcortical areas from the brains of patients with multiple sclerosis and control tissue from unaffected individuals was obtained from the web browser accompanying the original publication (https://cells.ucsc.edu/ms). AUCs were calculated by comparing cells from multiple sclerosis and control patients.

*Smillie et al., 2019*^69^. scRNA-seq data from colon biopsies of ulcerative colitis (UC) patients and healthy individuals was obtained from the Broad Institute Single Cell Portal (accession: SCP259). Cells from epithelial stromal, and immune fractions were combined into a single matrix, and AUCs were calculated by comparing cells from UC patients (including both inflamed and adjacent non-inflamed tissue) and controls.

*Wagner et al., 2018*^70^. scRNA-seq data from zebrafish embryos between 14-16 hours post-fertilization, with either the chordin locus or a control locus (tyrosinase) disrupted by CRISPR-Cas9 knockout, was obtained from GEO (accession: GSE112294). AUCs were calculated by comparing cells from chordin- or tyrosinase-targeted embryos.

*Wang et al., 2018*^15^. STARmap data from the visual cortex of mice housed in the dark and exposed to light for 0 h or 1 h before sacrifice was obtained from the STARmap website (https://www.starmapresources.com/data), and converted to count matrices using the STARmap python package (https://github.com/weallen/STARmap). Cells with less than 200 or more than 2000 counts were removed. The Rerg gene was removed from analysis as it was not measured in some samples. Cells with unassigned types were also removed, and the data was normalized to counts per million. AUCs were calculated by comparing cells from light-exposed and control mice.

*Wirka et al., 2019*^71^. scRNA-seq data from the aortic root of mice fed a high-fat diet or normal chow for eight weeks was obtained from GEO (accession: GSE131776). Metadata, including cell type annotations, was provided by the authors, and unannotated cells were removed. AUCs were calculated by comparing cells from high-fat diet and normal chow-fed mice.

*Wu et al., 2017*^55^. scRNA-seq data from the amygdala of mice subjected to 45 min of immobilization stress and control mice, and dissociated in the presence of actinomycin D following the Act-seq protocol, was obtained from GEO (accession: GSE103976). For RNA velocity analysis, raw sequencing data was obtained from the SRA (accession: PRJNA407818) and processed as described above. AUCs were calculated by comparing cells from stressed and control mice.

*Ximerakis et al., 2019*^72^. scRNA-seq data from whole brains of young (2-3 mo) and old (21-23 mo) mice was obtained from the Broad Institute Single Cell Portal (accession: SCP263), using coarse cell annotations as cell type labels. AUCs were calculated by comparing cells from young and old mice.

*Zhang et al., 2019*^30^. scRNA-seq data from synovial tissues from patients with rheumatoid arthritis (RA) or osteoarthritis (OA), obtained from ultrasound-guided biopsies or joint replacements, were obtained directly from the authors of the original publication. AUCs were calculated by comparing cells from RA and OA patients.

### Application of Augur to TESS

To experimentally validate the ability of Augur to uncover new biological mechanisms and identify neuron subtypes involved in complex behaviors, we applied Augur to investigate the neural circuits underlying the functional response to targeted epidural electrical stimulation (TESS) using single-nucleus transcriptomics.

#### Animal model

Experiments were conducted on adult male or female C57BL/6 mice (15-35 g body weight, 12-30 weeks of age). Vsx2:Cre (MMRRC//036672-UCD) transgenic mice were used and maintained on a mixed genetic background (129/C57BL/6). Housing, surgery, behavioral experiments and euthanasia were performed in compliance with the Swiss Veterinary Law guidelines. Animal care, including manual bladder voiding, was performed twice daily for the first 3 weeks after injury and once daily for the remaining post-injury period. All procedures and surgeries were approved by the Veterinary Office of the Canton of Geneva (Switzerland).

#### Surgical procedures and post-surgical care

Surgical procedures were performed as previously described^73–76^. Briefly, a laminectomy was made at the mid-thoracic level (T9 vertebra). We performed a contusion injury using a force-controlled spinal cord impactor (IH-0400 Impactor, Precision Systems and Instrumentation LLC, USA^77^), as previously described^73,78^. The applied force was set to 90 kdyn^73^. We next performed a partial laminotomy over spinal segments L2 and S1 and positioned the epidural stimulation electrodes (AS632, Cooner Wire, USA), and secured them at the midline to the dura. A common ground wire (∼1 cm of Teflon removed at the distal end) was inserted subcutaneously over the right shoulder. All wires were connected to a percutaneous amphenol connector (Omnetics Connector Corporation, USA) cemented to the skull of the animal. Analgesia (buprenorphine, Essex Chemie AG, Switzerland, 0.01–0.05 mg per kg, s.c.) was provided for three days after surgery.

#### Electrochemical stimulation

To reactivate lumbar motor circuits immediately prior to sacrifice and tissue harvest, we applied our previously described electrochemical neuromodulation therapy consisting of a serotoninergic replacement therapy and epidural electrical stimulation^73,75^. Briefly, five minutes before training, mice received a systemic (i.p.) administration of quipazine (5-HT2A/C, 0.3–0.6 mg/kg) and subcutaneous 8-OH-DPAT (5-HT1A/7, 0.1–0.2 mg/kg). Continuous epidural electrical stimulation (100 Hz bursts of 6 pulses at 0.2 ms, 100–300 *µ*A, 700 Hz) was delivered through L2 and S1 electrodes. This therapy immediately restored locomotion. Training was conducted quadrupedally on a three-dimensional robot with adjustable robotic body weight support against gravity (Robomedica, USA).

#### Kinematic recordings

Kinematics were recorded according to our previously described procedures, which have been extensively detailed^24,73–75,79,80^. Bilateral leg kinematics were captured using a Vicon high-speed motion capture system (Vicon Motion Systems, UK), consisting of 12 infrared cameras (200 Hz). We attached reflective markers bilaterally at the iliac crest, the greater trochanter (hip joint), the lateral condyle (knee joint), the lateral malleolus (ankle), and the distal end of the fifth metatarsophalangeal joint.

#### Kinematic analysis

For both the left and right legs, 15 step cycles were extracted for each mouse. A total of 75 parameters quantifying kinematic and kinetic features were computed for each leg and each gait cycle accordingly^24,73–75,79,80^. To evaluate differences between experimental conditions and groups, as well as the most relevant parameters to explain these differences, we implemented a multistep statistical procedure based on principal component analysis, as previously described^24,73–75,79,80^.

#### Single-nucleus RNA sequencing

Single nucleus dissociation was completed with a modified protocol based on our previous work^67^. Briefly, animals were euthanized by isoflurane inhalation and cervical dislocation. The thoracic SCI site was rapidly dissected and frozen on dry ice. Spinal cords were dounced in 500 *µ*l sucrose buffer (0.32 M sucrose, 10 mM HEPES [pH 8.0], 5 mM CaCl2, 3 mM Mg-acetate, 0.1 mM EDTA, 1 mM DTT) and 0.1% Triton X-100 with the Kontes Dounce Tissue Grinder. 2 mL of sucrose buffer was added and filtered through a *µ*m cell strainer. The lysate was subsequently centrifuged at 3200 g for 10 min at 4°C. The supernatant was decanted, and 3 mL of sucrose buffer added to the pellet and incubated for 1 min. The pellet was homogenized using an Ultra-Turrax and 12.5 mL of density buffer (1 M sucrose, 10 mM HEPES [pH 8.0], 3 mM Mg-acetate, 1 mM DTT) was added below the nuclei layer. The tube was centrifuged at 3200 g at 4°C and supernatant immediately poured off. The nuclei on the bottom half of the tube wall were collected with 100 *µ*l PBS with 0.04% BSA and 0.2 U/*µ*l RNase inhibitor. Resuspended nuclei were filtered through a 30 *µ*m strainer. The nuclei suspension was finally adjusted to 1000 nuclei/*µ*l.

#### Library preparation

Library preparation was carried out with 10x Genomics Chromium Single Cell Kit Version 2. The nuclei suspension was added to the Chromium RT mix to achieve loading numbers of 5,000. For downstream cDNA synthesis (13 PCR cycles), library preparation and sequencing, the manufacturer’s instructions were followed.

#### Read alignment

Reads were aligned to the latest Ensembl release (GRCm38.93), and a matrix of unique molecular identifier (UMI) counts was obtained using CellRanger count^81^. Velocyto^19^ was subsequently used to obtain count matrices of exonic and intronic reads. Seurat^44^ was used to calculate quality control metrics, including the number of genes detected, number of UMIs per cell, and % mitochondrial genes in order to filter low-quality cells appropriately (nUMI < 200; genes expressed in < 3 cells; % mitochondrial reads > 5%). The matrix used for downstream analysis consisted of 19,954 genes and 18,514 cells.

#### Clustering and integration

To integrate datasets across different experimental conditions, we took advantage of recently developed bioinformatic tools that align datasets from multiple conditions into a unified space^44^. Gene expression data was first normalized using regularized negative binomial models^82^, then integrated across batches using Seurat^44^. Batch effects were regressed out using the ‘latent.vars’ argument. Normalized and integrated gene expression matrices were clustered using Seurat^44^ to identify cell types in the integrated dataset using a standard workflow, including highly variable gene identification, principal component analysis, nearest-neighbor graph construction, and graph-based community detection. Following the identification of coarse-grained cell types (e.g., ‘neuron’), we identified fine-grained neuron subtypes by sub-clustering major cell types. We used clustering trees^83^ to guide the decision of the optimal resolution (**Supplementary Fig. 10a**). Cell types were manually annotated by using differential expression analysis to identify marker genes^6,44^. Putative cell types were assigned on the basis of marker gene expression, guided by previous work^67,84–86^.

#### RNA velocity

RNA velocity was calculated using the ‘velocyto’ R package^19^. Velocyto estimates cell velocities from their spliced and unspliced mRNA content. We generated the annotated spliced and unspliced reads using the ‘run10x’ function of the Velocyto command line tool, as described above. We then calculated gene-relative velocity using kNN pooling with k=10 (default).

#### Viral tract tracing

All surgeries on mice were performed at EPFL under general anaesthesia with isoflurane in oxygen-enriched air using an operating microscope, and rodent stereotaxic apparatus (David Kopf). To trace the efferent connections of Vsx2 (V2a) neurons AAV-DJ-hSyn Flex mGFP 2 A synaptophysin mRuby (Stanford Vector Core Facility, reference AAV DJ GVVC-AAV-100, titer 1.15E12 genome copies per ml^87^) was injected on each side of the cord of Vsx2-Cre mice at the L2 spinal level, 0.25 *µ*L 0.6 mm below the surface at 0.1 *µ*L per minute using glass micropipettes (ground to 50 to 100 *µ*m tips) connected via high-pressure tubing (Kopf) to 10-*µ*L syringes under the control of microinfusion pumps.

#### Immunohistochemistry

After terminal anaesthesia by barbiturate overdose, mice were perfused transcardially with 4% paraformaldehyde and spinal cords processed for immunofluorescence as previously described^73,88^. Primary antibodies were: goat anti-choline acetyltransferase (ChAT, 1:50, Millipore, AB144P). Secondary antibodies were: Alexa Fluor 647 Donkey Anti Goat (1:200; Life Technologies, AB32849). Immunofluorescence was imaged digitally using a slide scanner [Olympus VS-120 Slide scanner] or confocal microscope [Zeiss LSM880 + Airy fast module with ZEN 2 Black software (Zeiss, Oberkochen, Germany)]. Images were digitally processed using ImageJ (NIH) or Imaris (Bitplane, v.9.0.0).

#### RNAscope

We confirmed the *in situ* localization of cell type markers and the expression of the immediate early gene Fos using RNAscope. Briefly, 16 *µ*m cryosections were obtained from fixed-frozen spinal cords of animals undergoing identical experimental procedures. We used these sections to confirm the localization of Spp1 (cat. no. 435191), Slc6a5 (cat. no. 409741-C3) and Vsx2 (cat. no. 438341). We additionally included an analysis of negative controls that were not prioritized by Augur including Cck (cat. no. 402271-C3), Npy (cat. no. 313321), Rorb (cat. no. 444271-C3), Pnoc (cat. no. 437881), Gal (cat. no. 400961-C3), and Trh (cat. no. 436811 neurons). These cell types have also been validated elsewhere^67,84–86^. We combined gene markers with Fos (cat. no. 316921-C2) to confirm the presence of immediate early gene activation in these cell types^67^. To detect the transcripts we used the RNAscope assay for fixed frozen tissue (Advanced Cell Diagnostics)^89^. Probes were designed and provided by Advanced Cell Diagnostics, Inc. Staining was performed according to standard procedures, using the RNAscope Fluorescent Multiplex Reagent Kit (cat. no. 323133).

### Visualization

Throughout the manuscript, box plots show the median (horizontal line), interquartile range (hinges) and smallest and largest values no more than 1.5 times the interquartile range (whiskers), and error bars show the standard deviation.

## Supporting information

Supplementary Software

## Code availability

Augur is available from GitHub (https://github.com/neurorestore/Augur) and as **Supplementary Software 1**.

## Data availability

Raw sequencing data and count matrices have been deposited to the Gene Expression Omnibus.

## Acknowledgements

We thank D. Arneson, D. Avey, R. Mitra, A. Haber, O. Yilmaz, G. Chew, J. Polo, L. Adlung, I. Amit, D.W. Kim, D.J. Anderson, M. Basiri, R. Wirka, T. Quertermous, and F. Zhang for providing data and/or cell type annotations. This work was supported by a Consolidator Grant from the European Research Council [ERC-2015-CoG HOW2WALKAGAIN 682999] (to G.C.), the Swiss National Science Foundation (subside 310030_185214, to G.C.), Genome Canada and Genome British Columbia (project 214PRO, to L.J.F.), and Wings for Life (to M.A.S.). This work was also supported in part by the Intramural Research Program of the NIH, NINDS (to K.M. and A.L.). This work was enabled in part by the support provided by WestGrid and Compute Canada (to A.A.P. and L.J.F.). M.A.S. is supported by a CIHR Vanier Canada Graduate Scholarship, an Izaak Walton Killam Memorial Pre-Doctoral Fellowship, a UBC Four Year Fellowship, and a Vancouver Coastal Health–CIHR–UBC MD/PhD Studentship, and acknowledges travel support from a Brain Canada Hubert van Tol fellowship and a BCRegMed Collaborative Research Travel Grant. J.W.S. is supported by a CIHR Banting Postdoctoral fellowship, an Alberta Innovates postdoctoral fellowship, and a Killam Postdoctoral fellowship.

## Author contributions

M.A.S. and J.W.S. contributed equally to this work. M.A.S. and J.W.S. designed and implemented Augur, and performed all computational analyses. M.A.S., J.W.S., and M.G. processed published datasets. J.W.S., C.K., M.A.A., T.H.H., and M.M. performed experimental validation work, including viral tract tracing and RNAscope. C.K., K.J.E.M., and A.J.L. performed nucleus extraction and single-nucleus RNA-seq. M.G. and Q.B. analyzed experimental validation data. A.A.P., L.F., G.L.M., and G.C. supervised the work. M.A.S., J.W.S., and G.C. wrote the manuscript. All authors contributed to its editing.

---

**Supplementary Figure 1.**
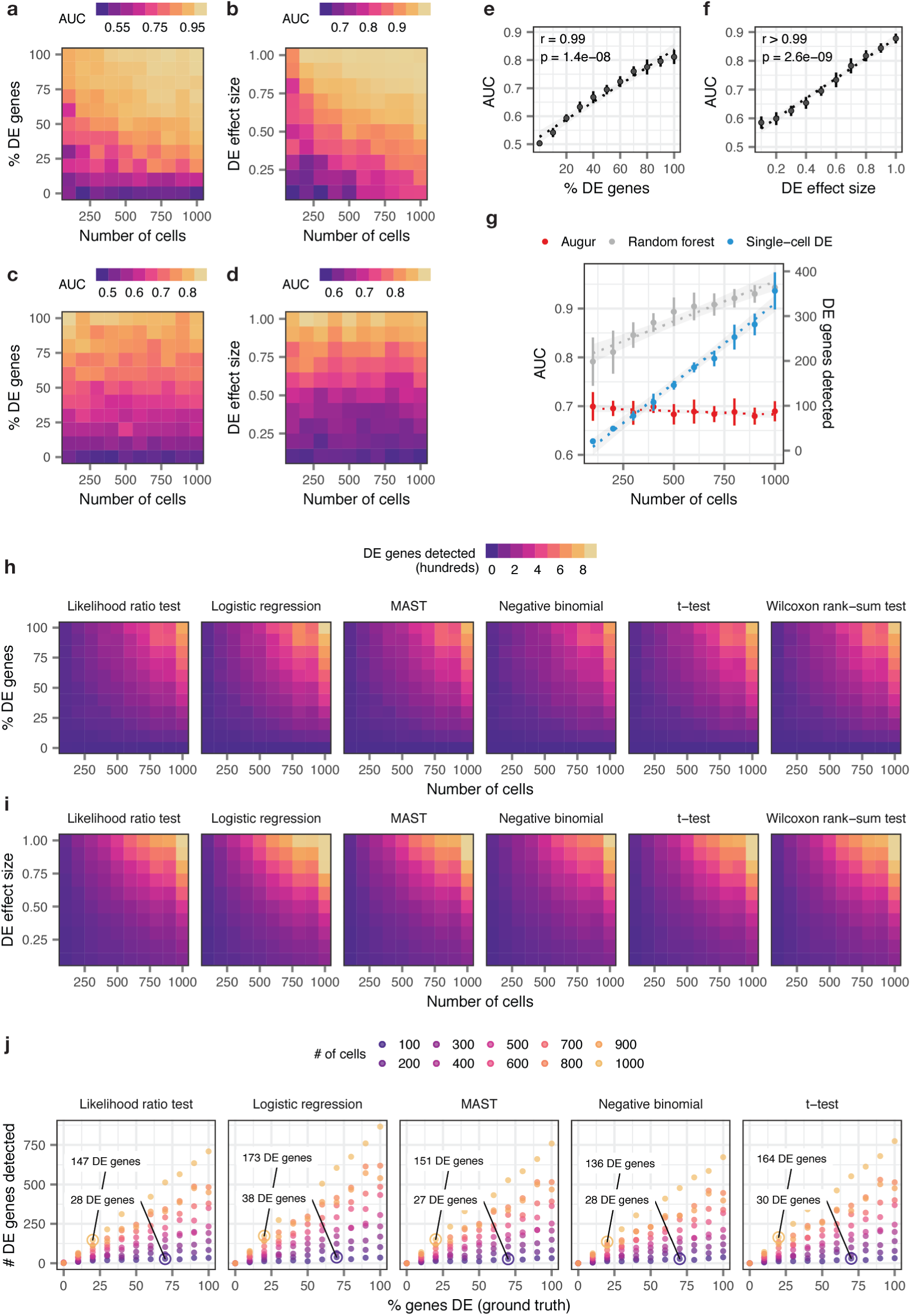
Augur overcomes confounding factors to cell type prioritization in simulated single-cell RNA-seq datasets. **a-b**, Area under the receiver operating characteristic curve (AUC) of a random forest classifier trained in three-fold cross-validation to distinguish two simulated populations of cells^49^, with the total number of cells increasing from 100 to 1,000 and the proportion of differentially expressed genes between the two populations varying from 0% to 100%, **a**, or the location parameter of the differential expression factor log-normal distribution varying from 0.1 to 1.0, **b**. **c-d**, As in **a-b**, but with the naive random forest classifier replaced with the subsampling procedure employed by Augur. **e-f**, Relationship between Augur AUC and the proportion of differentially expressed genes, **e**, or the location parameter of the differential expression factor log-normal distribution, **f**, in distinguishing two simulated populations (*n* = 200 cells total). The mean and standard deviation of 10 simulation replicates are shown. **g**, Cell type prioritizations (AUC or number of differentially expressed genes) for a naive random forest classifier, Augur, and an exemplary single-cell differential expression test^6^, the Wilcoxon rank-sum test, for two simulated populations of cells with 50% of genes differentially expressed and a log-normal location parameter of 0.5, with the total number of cells increasing from 100 and 1,000 cells. Like a naive random forest strategy, the number of differentially expressed genes detected by the Wilcoxon rank-sum test scales linearly with the number of cells. **h-i**, Number of differentially expressed genes detected by six tests for single-cell differential gene expression between two simulated populations of cells, with the total number of cells increasing from 100 to 1,000 and the proportion of differentially expressed genes between the two populations varying from 0% to 100%, **h**, or the location parameter of the differential expression factor log-normal distribution varying from 0.1 to 1.0, **i**. **j**, Relationship between number of differentially expressed genes detected by five tests for single-cell differential gene expression and the proportion of differentially expressed genes simulated between the two populations, for simulated populations of between 100 and 1,000 cells (see also **Fig. 1e**). All single-cell differential expression tests detect a larger number of differentially expressed genes in a large population of cells with modest transcriptional perturbation (20% of genes differentially expressed) than in a smaller population of cells with more profound perturbation (70% of genes differentially expressed).

**Supplementary Figure 2.**
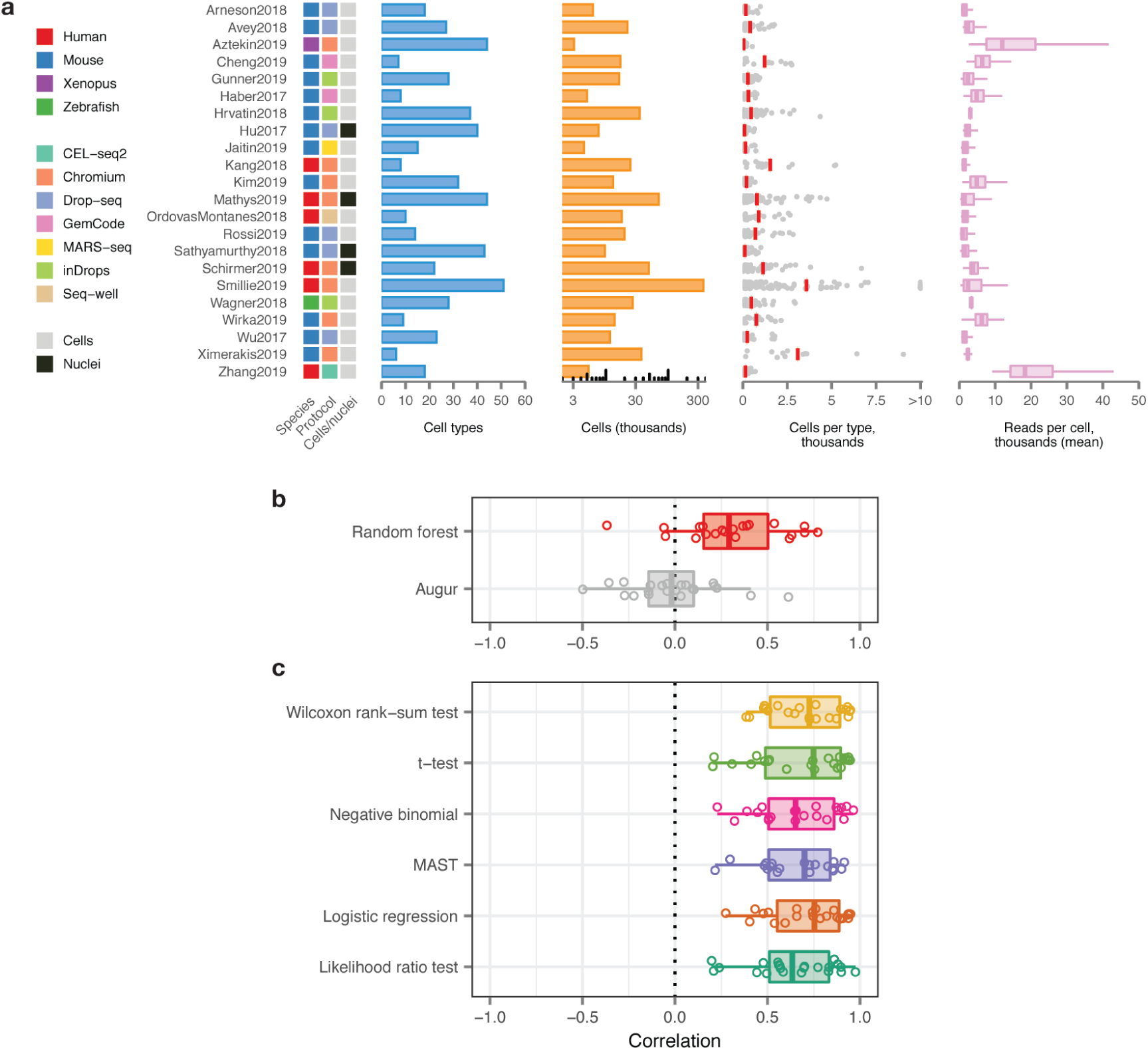
Augur overcomes confounding factors to cell type prioritization in a compendium of published single-cell RNA-seq datasets. **a**, Overview of 22 published scRNA-seq datasets comparing two or more experimental conditions, used to verify the relationship between cell type prioritizations from a random forest classifier, Augur, or single-cell differential expression tests. Left, heatmap indicating the species of origin, the sequencing protocol, and whether cells or nuclei were sequenced. Right, properties of each dataset, including the total number of cell types identified in the original studies; the total number of cells sequenced; the number of cells per type (red bars indicate mean); and the mean number of reads for cells of each type. **b**, Pearson correlations between the AUC of each cell type, and the number of cells of that type sequenced, across 22 datasets for Augur, bottom, and a naive random forest classifier without subsampling, top, as shown in **Fig. 2c**. **c**, Pearson correlations between the number of differentially expressed genes per cell type, at 5% FDR, and the number of cells of that type, sequenced across 22 datasets for six statistical tests for differential expression.

**Supplementary Figure 3.**
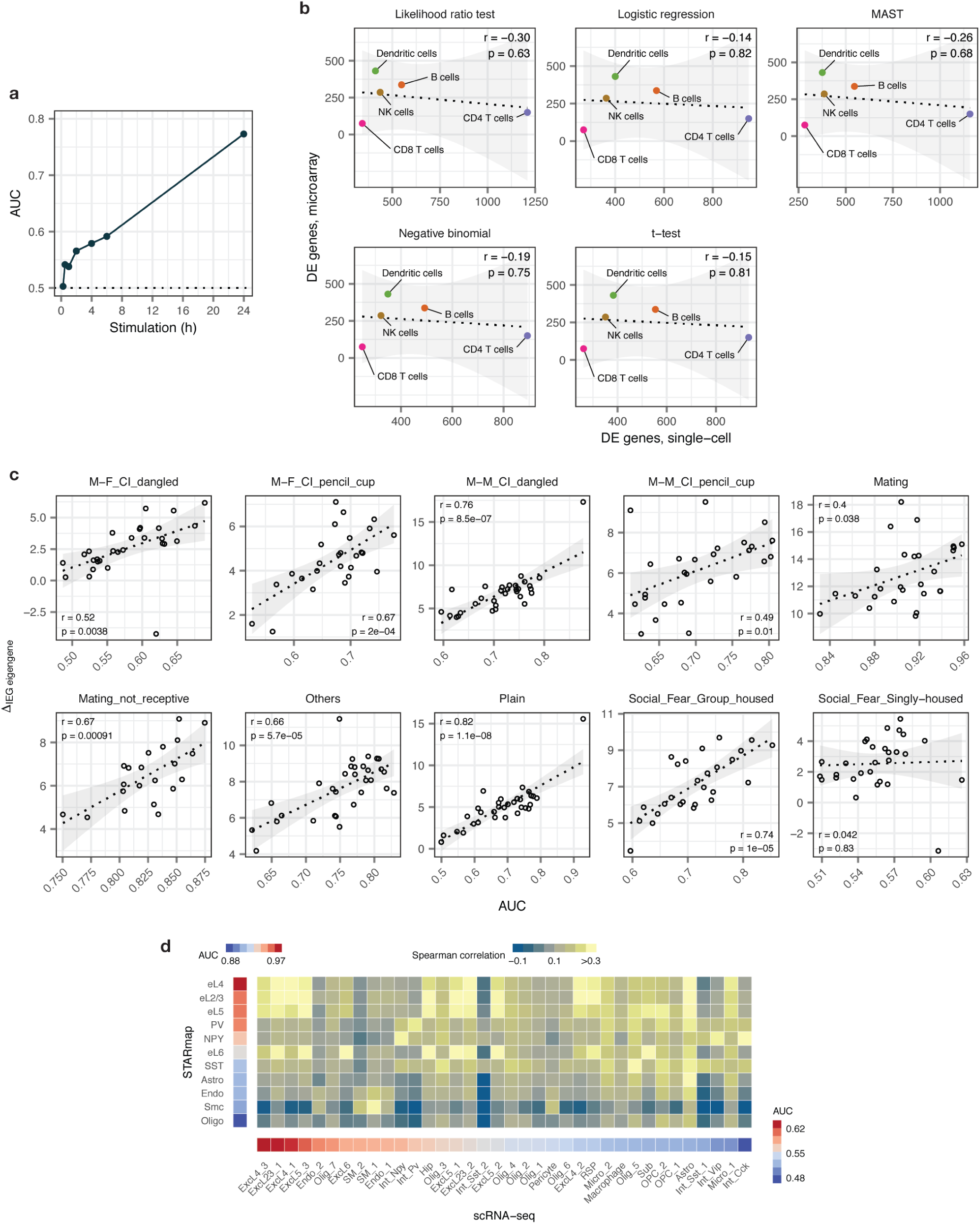
Evaluation of cell type prioritization across single-cell modalities. **a**, AUCs of Jurkat cells stimulated with PMA/ionomycin for seven time points between 15 min and 24 h, compared to control cells^13^. Augur detects the expected dose-response relationship of T cell activation. **b**, Relationship between the number of differentially expressed genes by microarray profiling of FACS-sorted mouse PBMCs stimulated with interferon for 2 h^8^, and the number of differentially expressed genes in an independent single-cell RNA-seq study of human PBMCs stimulated with interferon for 6 h^4^, detected by five statistical tests for single-cell differential gene expression. **c**, Relationship between AUCs for cell types of the ventromedial hypothalamus of mice exposed to one of ten behavioral stimuli, as computed by Augur, and the mean change in the first principle component of intermediate early gene (IEG) expression (ΔIEG eigengene), reflecting activation of IEG transcription in that cell type in response to the behavioral stimulus^9^. Augur cell type prioritizations reflect induction of neuronal intermediate early genes. See also **Fig. 1i**. **d**, Comparison of cell type prioritization in independent scRNA-seq and single cell imaging transcriptomics (STARmap) studies of the mouse visual cortex after light exposure. Left, Augur cell type prioritization in the STARmap dataset^15^. Bottom, Augur cell type prioritization in the scRNA-seq dataset^14^. Center, correspondence between cell types defined in the scRNA-seq and STARmap datasets, quantified as the Spearman correlation coefficient between average profiles for each cell type across 139 genes present in both datasets.

**Supplementary Figure 4.**
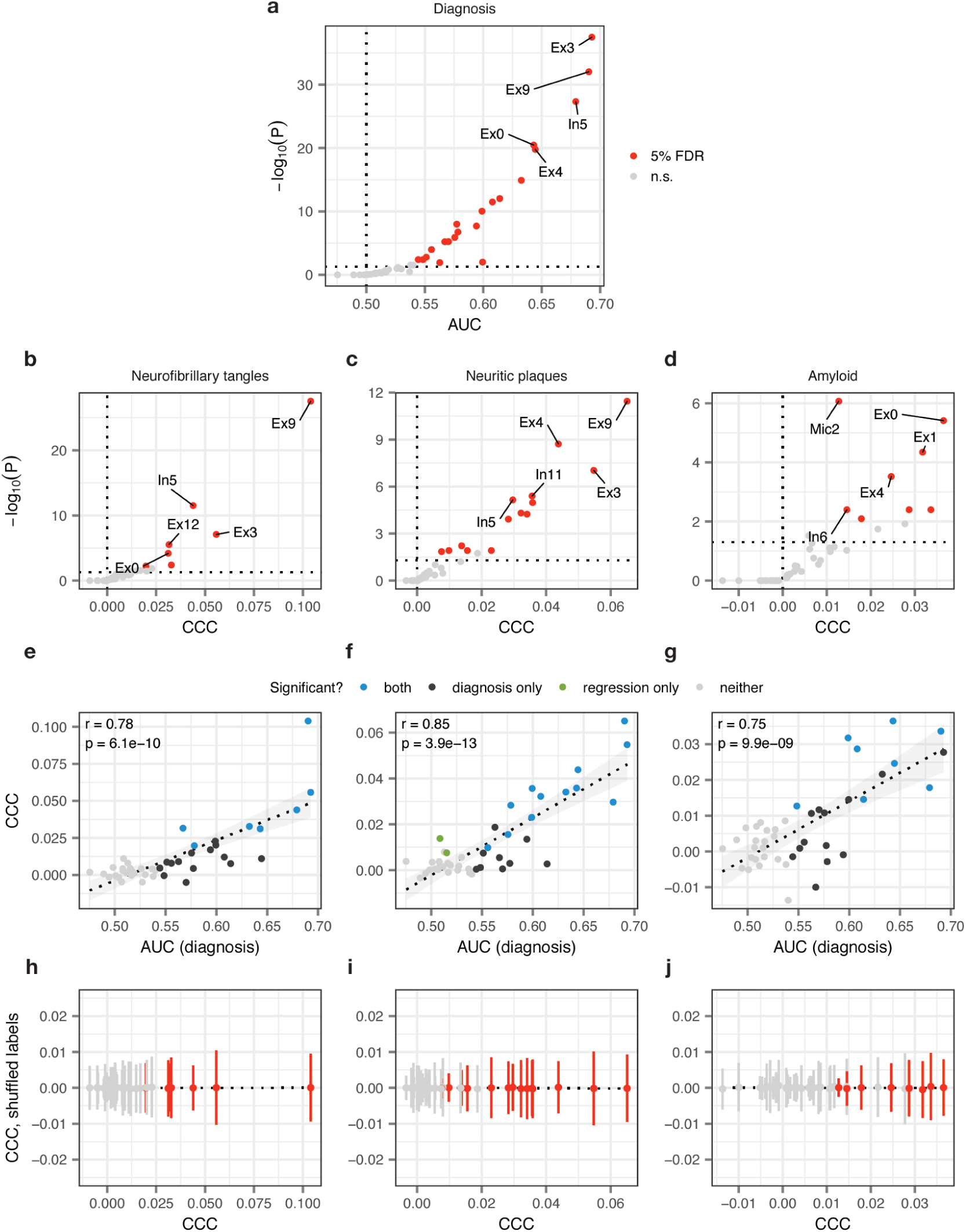
Cell type prioritization with continuous sample labels by random forest regression. **a**, Cell type prioritization (AUC) in post-mortem prefrontal cortex of individuals with Alzheimer’s disease and healthy controls in the Mathys et al., 2019 dataset^5^, and its statistical significance (permutation test; Methods). **b-d**, Cell type prioritization (concordance correlation coefficient, CCC) in the Mathys et al., 2019 dataset^5^ using continuous neuropathological variables as sample labels. **b**, Neurofibrillary tangle burden, as determined by microscopic examination of silver-stained slides from five brain regions. **c**, Neuritic plaque burden, as determined by microscopic examination of silver-stained slides from five brain regions. **d**, Overall amyloid-*β* level, as determined by immunohistochemistry of eight brain regions. **e-g**, Relationship between cell type prioritization in binary classification of Alzheimer’s disease vs. control individuals and regression of neurofibrillary tangle burden, **e**; neuritic plaque burden, **f**; and amyloid burden, **g**. Cell type prioritizations with continuous sample labels are strongly and significantly correlated with the case-control binary classification task. **h-j**, Cell type prioritization with shuffled sample labels for continuous outcome variables (neurofibrillary tangle burden, **h**; neuritic plaque burden, **i**; and amyloid burden, **j**). With shuffled labels, the CCC converges to zero.

**Supplementary Figure 5.**
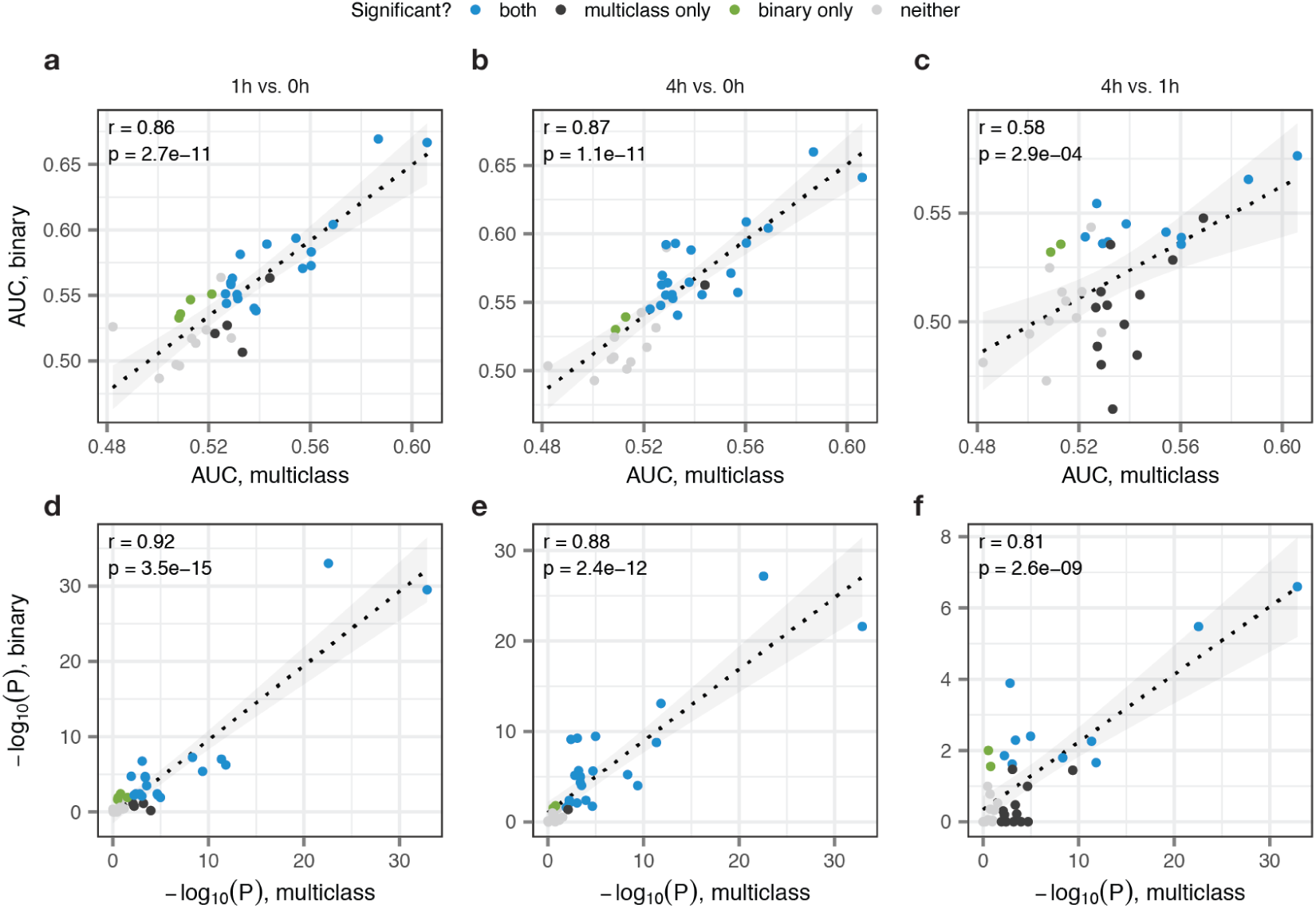
Cell type prioritization with three or more experimental conditions by multiclass classification. Comparison of binary and multiclass classification for cells of the mouse visual cortex after exposure to light in the Hrvatin et al., 2018 dataset^14^. **a-c**, Relationship between cell type prioritization in all possible pairwise binary classification tasks (**a**, 1 h vs. 0 h; **b**, 4 h vs. 0 h; **c**, 4 h vs. 1 h), and the corresponding multiclass classification task (including all three timepoints; macro-averaged AUC). Cell type prioritizations in multiclass classification are strongly and significantly correlated with each possible binary classification task, reflecting the incorporation of information from all three experimental conditions. **d-f**, Relationship between statistical significance of cell type prioritization in pairwise binary classification and multiclass classification, as assessed by permutation test (Methods). Cell types with the most significant transcriptional responses to light exposure are consistently detected in both binary and multiclass classification tasks.

**Supplementary Figure 6.**
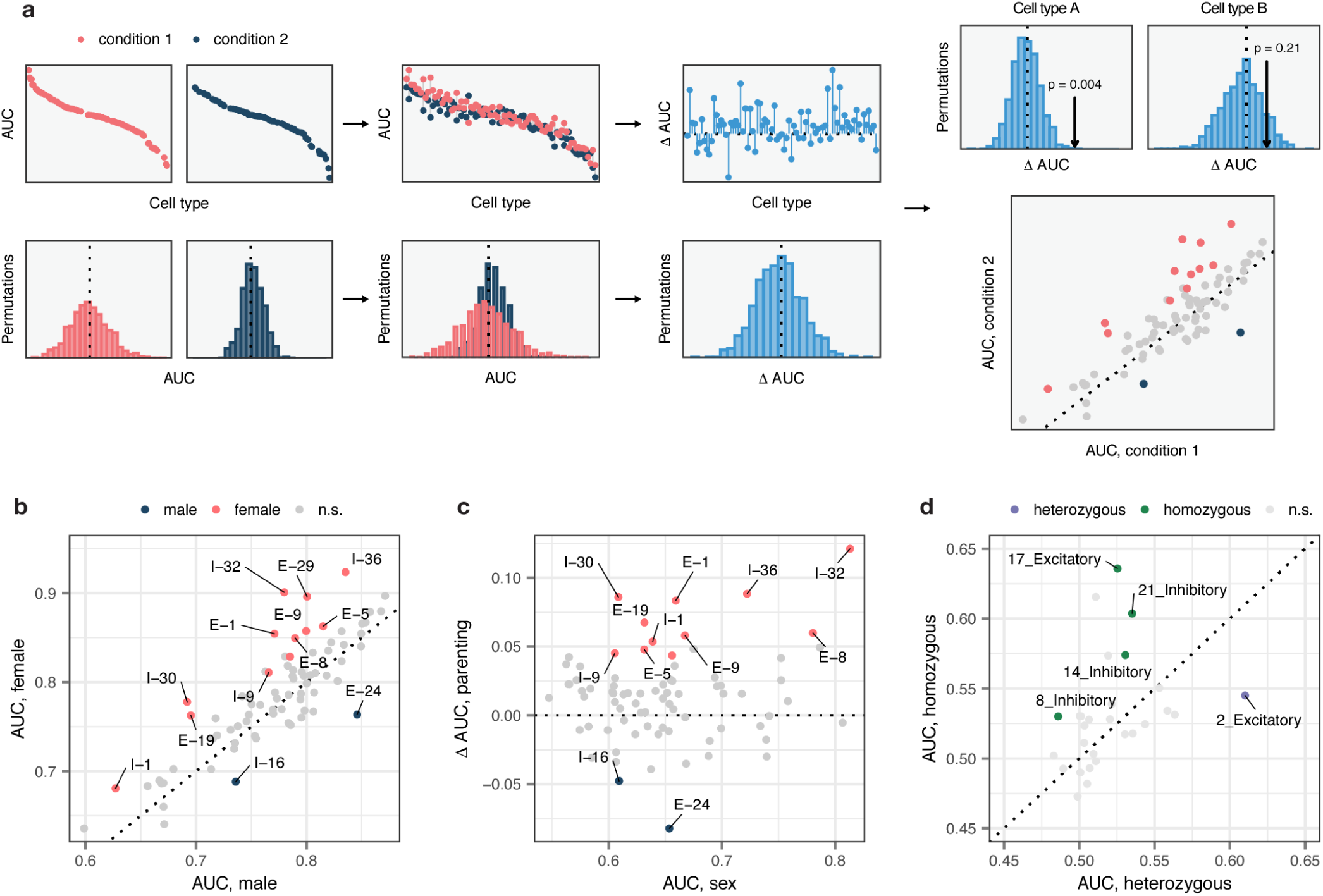
Differential cell type prioritization in single-cell RNA-seq data. **a**, Schematic overview of the permutation-based test for differential prioritization with Augur. First, cell type prioritization is performed within each of two conditions separately, yielding condition-specific AUCs for each cell type. Next, sample labels are randomly permuted within each cell type, and cell type prioritization is performed on shuffled data, yielding a null distribution of AUCs for each cell type and condition. AUCs for matching cell types are compared across conditions to calculate a ‘ΔAUC score’ for each cell type, and a null distribution of ΔAUC scores is calculated using the permuted data. Permutation p-values can then be calculated for each cell type, enabling the identification of statistically significant differences in cell type prioritization between conditions, as well as the condition in which the cell type is more transcriptionally separable. **b**, Neuron subtypes with statistically significant differences in AUC between female and male mice during parenting, in a single-cell imaging transcriptomics experiment employing multiplexed error robust fluorescence in situ hybridization (MERFISH)^17^. Eleven subtypes have significantly higher AUCs in female parents, whereas two have significantly higher AUCs in male parents. **c**, Relationship between differential prioritization ΔAUC for parenting between male and female mice, and AUC for sex in naive mice. Several neuronal subtypes preferentially activated during parenting in female mice are also transcriptionally distinct in naive mice, such as the I-32 cluster, which is enriched for aromatase expression, and expresses multiple sex steroid hormone receptors^17^. **d**, Neuron subtypes with statistically significant differences in AUC in response to whisker lesioning in Cx3cr1^+/−^ as compared to Cx3cr1^−/−^ mice, in a single-cell RNA-seq experiment^18^. Four subtypes are have significantly higher AUCs in homozygous mice, whereas one subtype has a significantly higher AUC in heterozygous mice.

**Supplementary Figure 7.**
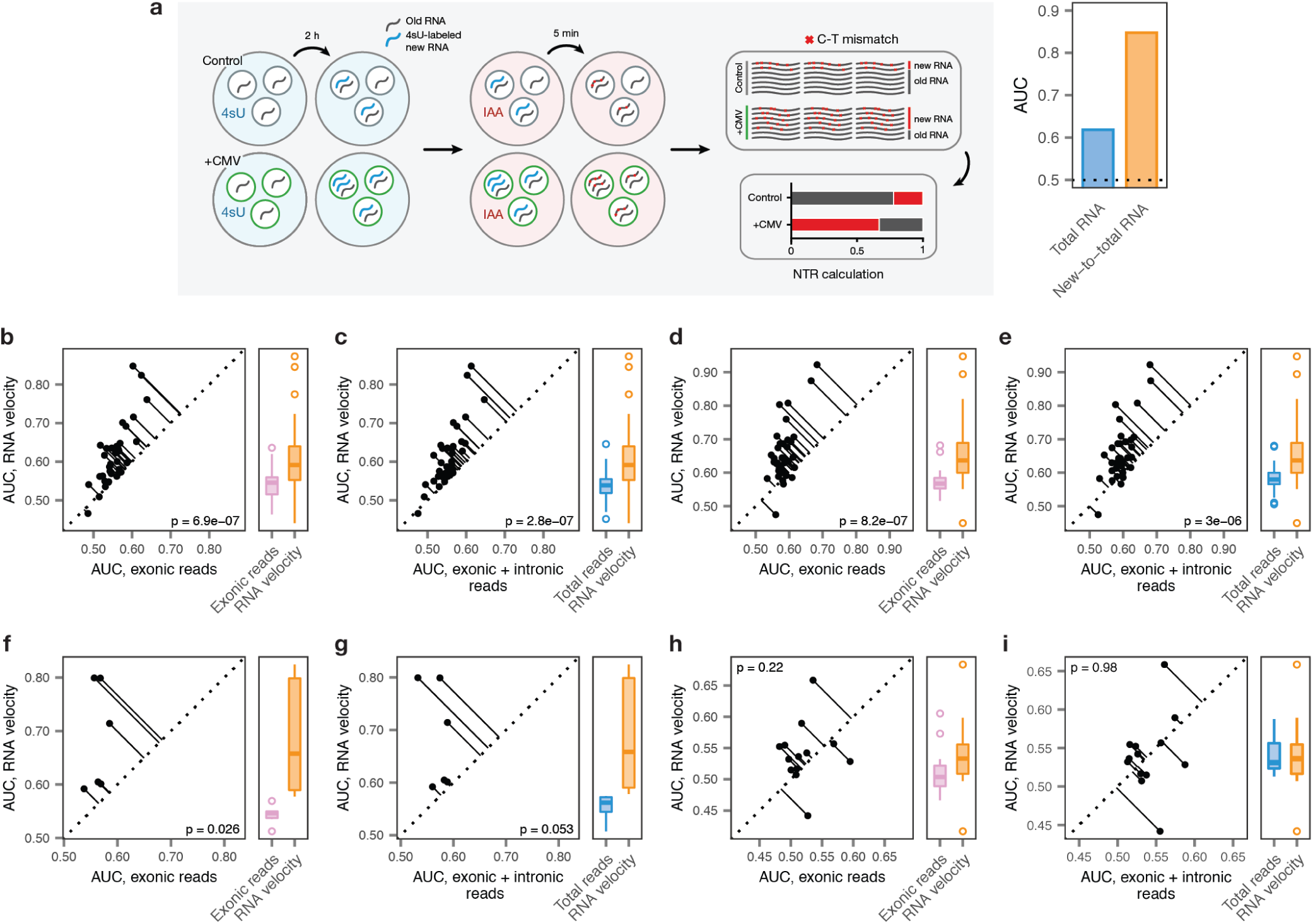
Cell type prioritization from transcriptional dynamics in acute experimental perturbations. **a**, Left, schematic overview of the scSLAM-seq^20^ workflow. Cells are exposed to the nucleoside analogue 4-thiouridine (4sU), which is incorporated during transcription and converted to a cytosine analogue by iodoacetamide prior to RNA sequencing. This labeling permits *in silico* deconvolution of RNA molecules transcribed before and after 4sU exposure (‘old’ and ‘new’, respectively), and calculation of the ratio of new to total RNA (NTR), an experimental analogue to the computationally determined ‘RNA velocity’^19,20^. Right, AUCs for mouse fibroblasts exposed to lytic mouse cytomegalovirus (CMV) at 2 h post-infection, calculated by applying Augur to either total RNA or the NTR. The greater separability for the NTR reflects additional information specifically captured by the temporal dynamics of RNA expression in the context of this acute perturbation^20^. **b-e**, Cell type prioritization based on exonic reads, total RNA, or RNA velocity for cells of the mouse visual cortex after exposure to light for 1 h, **b-c**, or 4 h, **d-e**, in the Hrvatin et al., 2018 dataset^14^. The AUC is significantly higher for RNA velocity than for either exonic reads (paired t-tests: **b**, 1 h, p = 6.9 × 10^−7^; **d**, 4 h, p = 8.2 × 10^−7^) or total RNA (**c**, 1 h, p = 2.8 × 10^−7^; **e**, 4 h, p = 3.0 × 10^−6^), reflecting additional information specifically captured by acute transcriptional dynamics. **f-g**, Cell type prioritization based on exonic reads, total RNA, or RNA velocity in an Act-seq^55^ dataset, which minimizes transcriptional changes induced by single-cell dissociation. Cell types of the medial amygdala in mice subjected to 45 min of immobilization stress and control mice were profiled by Drop-seq^56^ after treatment with the transcription inhibitor actinomycin D. The AUC is higher for RNA velocity than for either exonic reads (**f**, p = 0.026) or total RNA (**g**, p = 0.053), reflecting additional information specifically captured by acute transcriptional dynamics, and indicating this is not an artefact related to the transcriptional perturbations known to be induced by conventional dissociation procedures^90^. **h-i**, Cell type prioritization based on exonic reads, total RNA, or RNA velocity in a chronic perturbation. Cell types of the lateral hypothalamic area were profiled by Drop-seq^56^ in mice after 9-16 weeks of maintenance on either high-fat diet or control diet^12^. No significant difference in AUCs was observed for RNA velocity compared to either exonic reads (**h**, p = 0.22) or total RNA (**i**, p = 0.98), consistent with the time scale of the experimental perturbation.

**Supplementary Figure 8.**
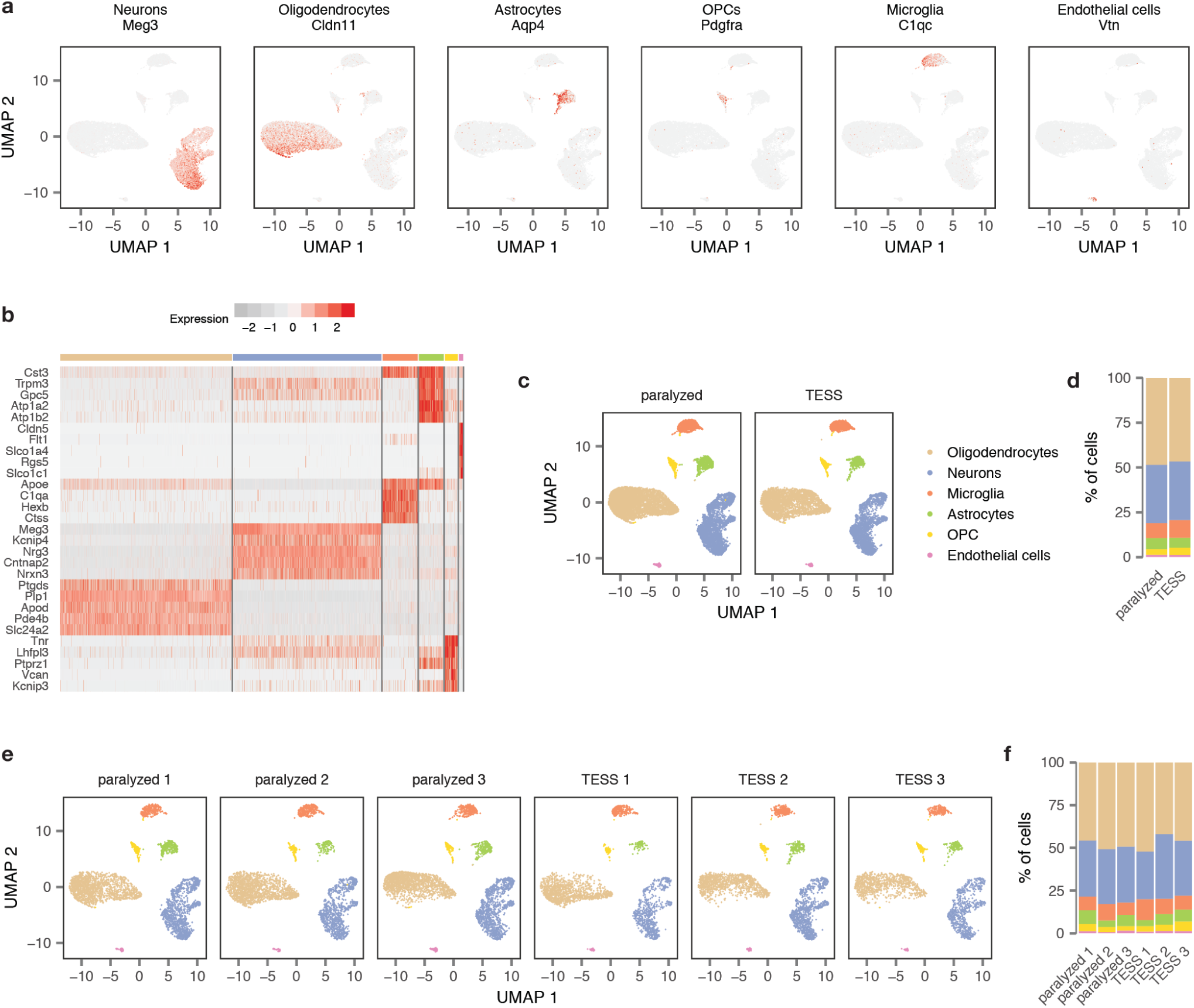
Single-nucleus RNA-seq detects the major cell types of the lumbar spinal cord across experimental conditions and replicates. **a**, Expression of key marker genes for the six major cell types of the lumbar spinal cord. **b**, Top five marker genes for each major cell type of the lumbar spinal cord. **c**, Cell type detection across experimental conditions. TESS, targeted electrical epidural stimulation of the lumbar spinal cord. **d**, Proportion of cells of each type detected in each experimental condition. **e**, Cell type detection across experimental replicates (*n* = 3 mice per condition). **f**, Proportion of cells of each type detected in each experimental replicate.

**Supplementary Figure 9.**
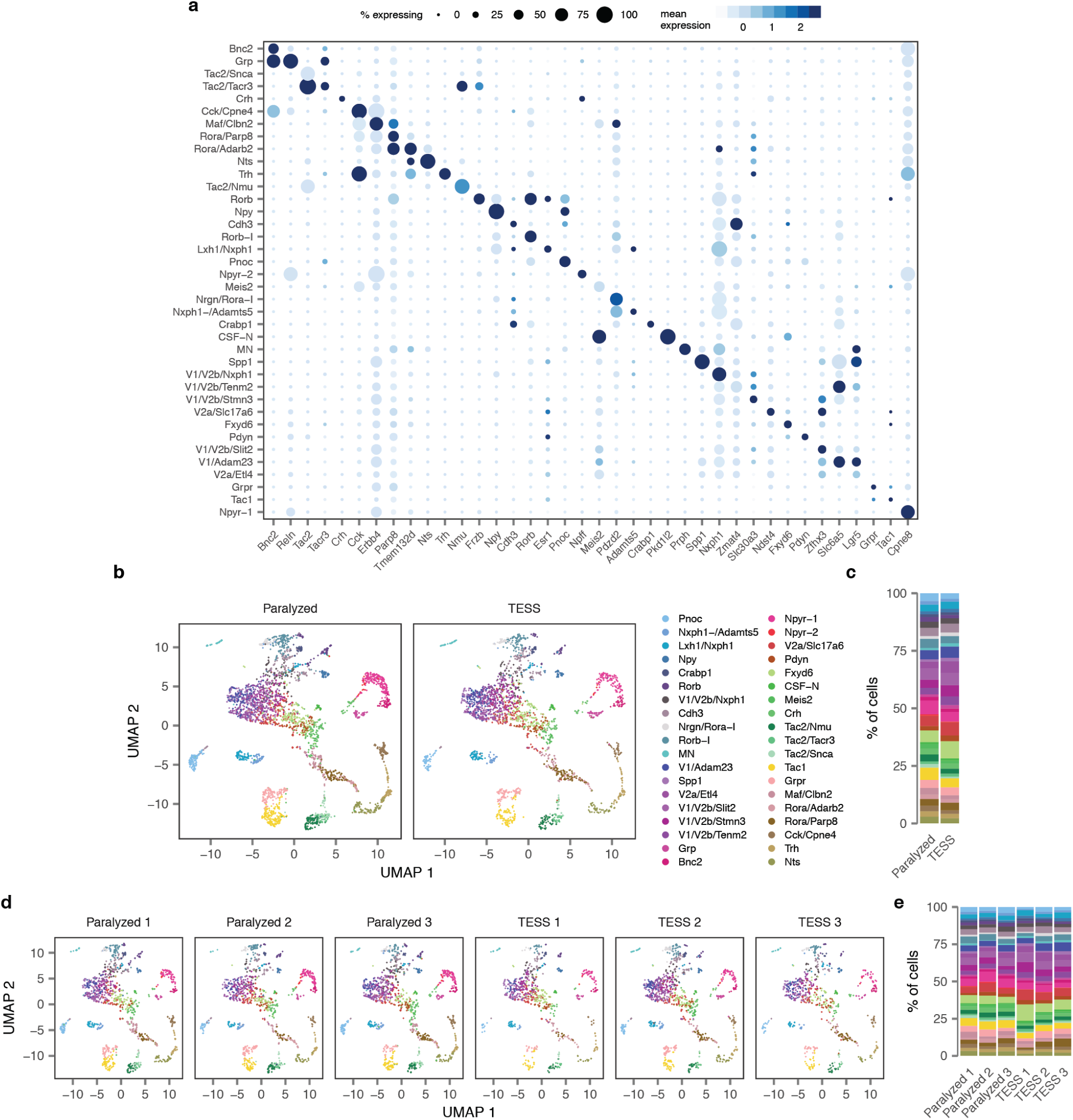
Subclustering of single-neuron transcriptomes identifies 39 neuron subtypes in the mouse lumbar spinal cord. See also **Supplementary Figure 10a**. **a**, Dot plot showing expression of one marker gene per cell type for the 39 neuron subtypes of the mouse lumbar spinal cord. **b**, Neuron subtype detection across experimental conditions. TESS, targeted electrical epidural stimulation of the lumbar spinal cord. **c**, Proportion of neurons of each subtype detected in each experimental condition. **d**, Neuron subtype detection across experimental replicates (*n* = 3 mice per condition). **e**, Proportion of neurons of each subtype detected in each experimental replicate.

**Supplementary Figure 10.**
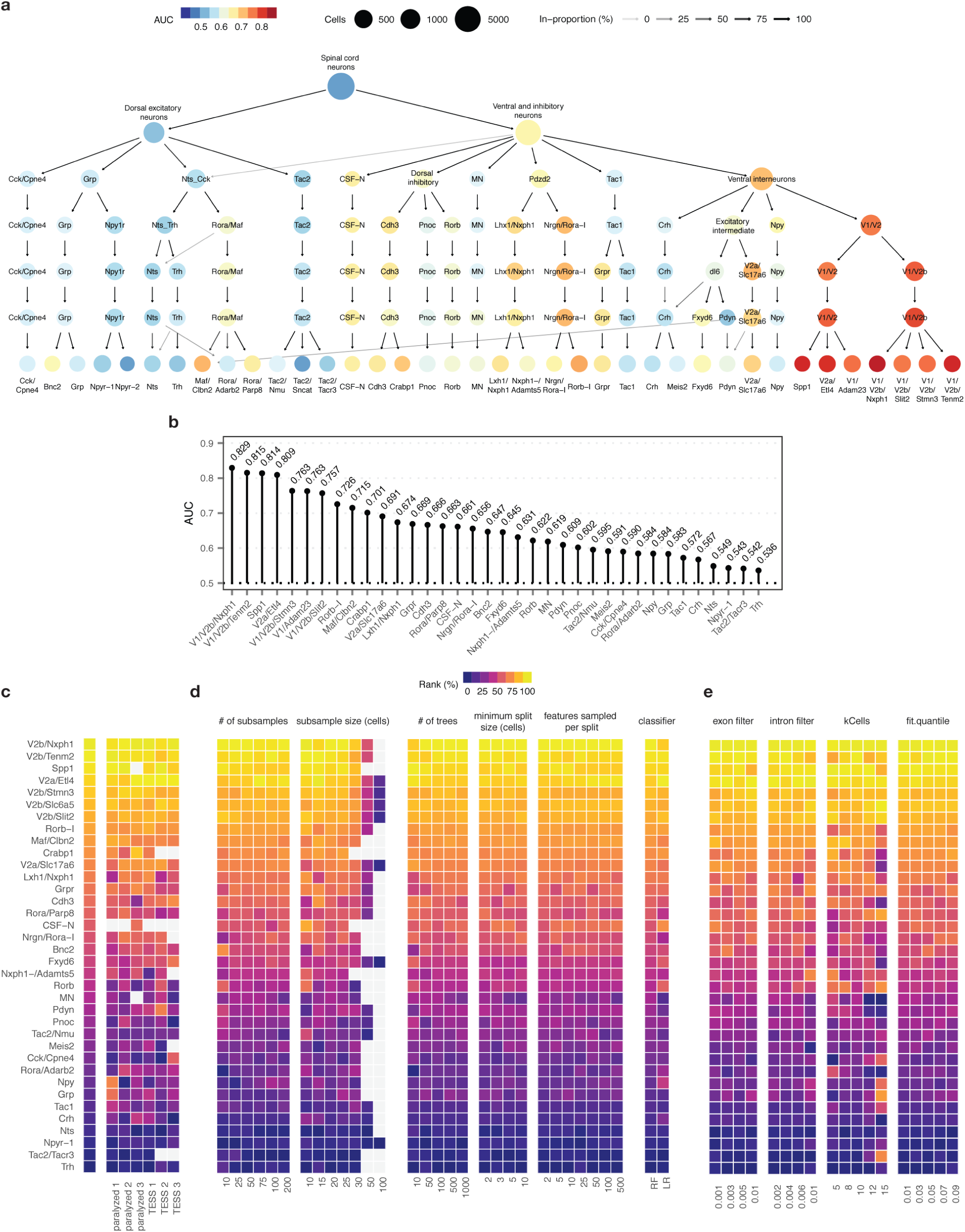
Robustness of Augur cell type prioritizations for mouse lumbar spinal cord neurons. **a**, Clustering tree^83^ of mouse spinal cord neurons over seven clustering resolutions, revealing the hierarchical relationships between spinal cord neuron subtypes. Node color reflects AUCs for cell type prioritization in targeted electrical epidural stimulation. **b**, AUCs for each of 37 neuron subtypes represented by at least 20 cells in both control and TESS-treated mice. **c-e**, Robustness of cell type prioritization for neuron subtypes of the mouse lumbar spinal cord. **c**, Impact of systematically withholding cells from each of six replicates (*n* = 3 per group) on cell type prioritization. Left, cell type prioritization with all six replicates, as in **Fig. 2f**. Grey tiles indicate neuron subtypes that were not represented by at least 20 cells in each condition after removal of cells from an experimental replicate. **d**, Impact of varying Augur parameters, including the number of subsamples and the size of each subsample; random forest-specific hyperparameters (number of trees, minimum split size, number of features sampled per split); and the choice of classifier (random forest, RF; L1-penalized logistic regression, LR) on cell type prioritization. Grey tiles indicate sample sizes larger than the number of cells of that type in the dataset. **e**, Impact of varying RNA velocity parameters, including exonic and intronic expression filters, the number of cells in the k-nearest neighbors pooling, and the extreme quantiles used to fit *γ* coefficients, on cell type prioritization.

**Supplementary Figure 11.**
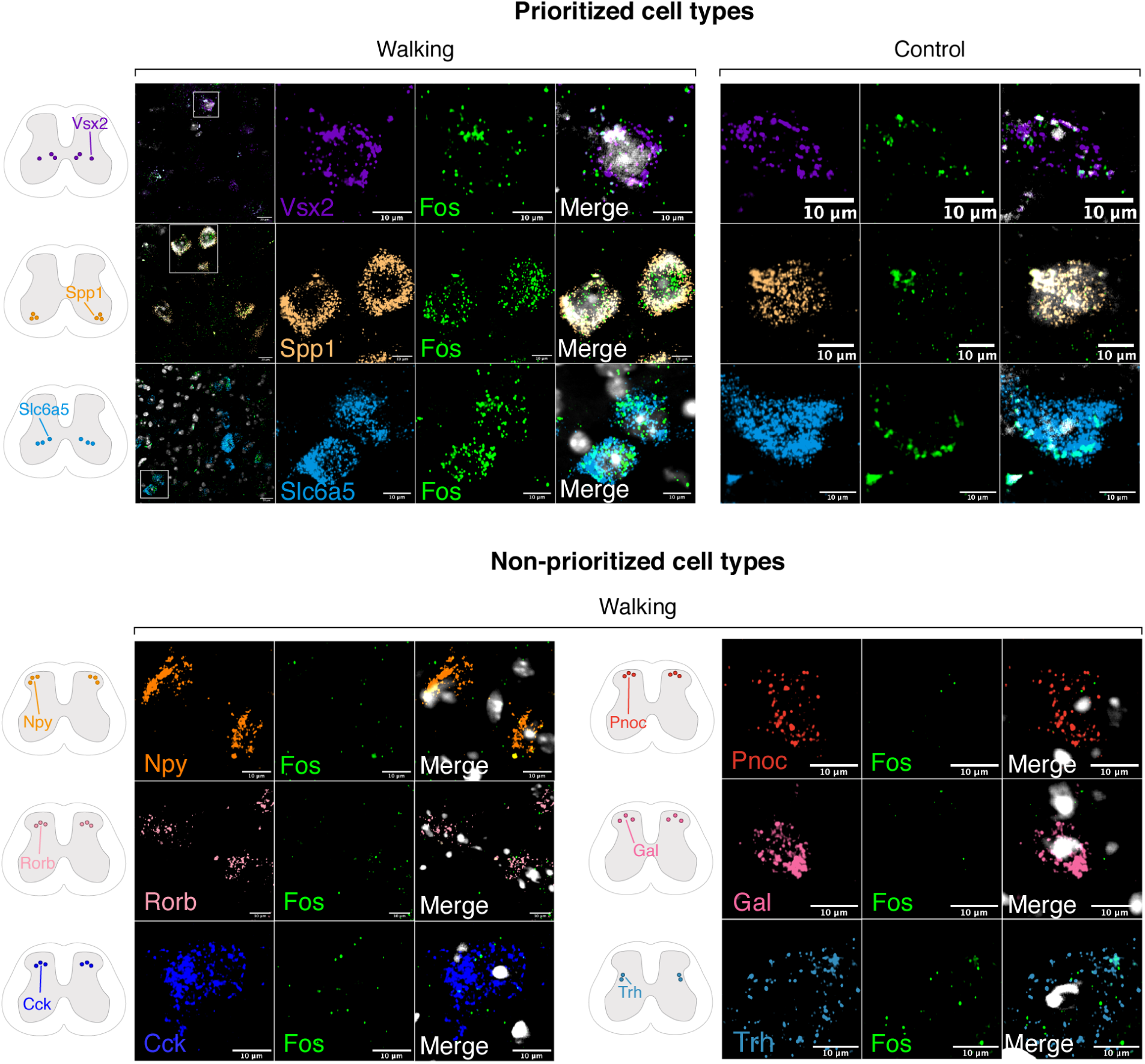
Absence of colocalization of canonical marker genes for cell types not prioritized by Augur and Fos by RNAscope *in situ* hybridization. Schematic indicates imaging location for each marker within the spinal cord.

**Supplementary Figure 12.**
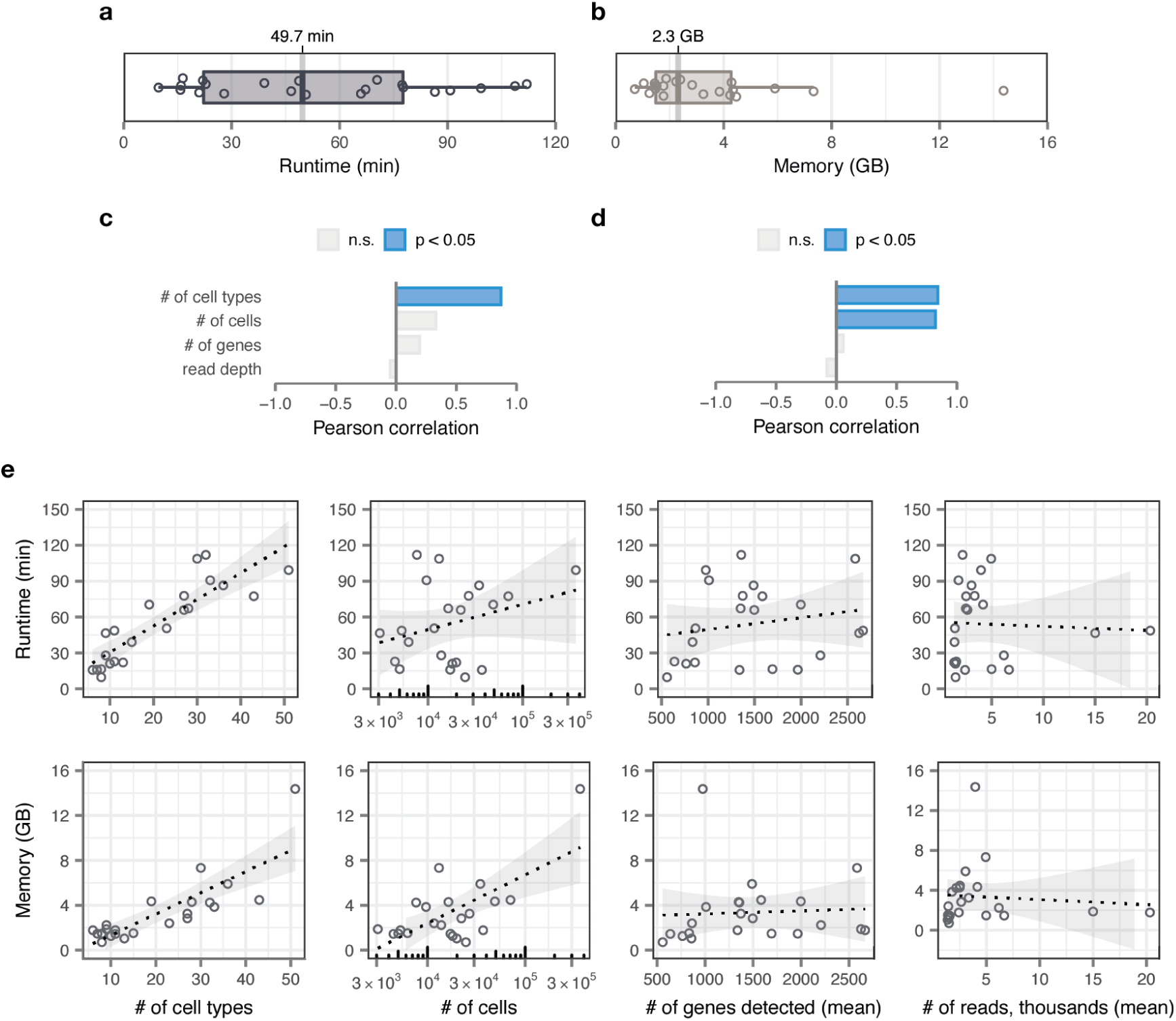
Computational efficiency of cell type prioritization. **a**, Elapsed real time (“wall time”) for Augur cell type prioritization on a compendium of 22 scRNA-seq datasets, with Augur run on four cores. The median wall time, 49.7 min, is highlighted. **b**, Peak memory usage for Augur cell type prioritization on the scRNA-seq dataset compendium. The median peak memory usage, 2.3 GB, is highlighted. **c-d**, Pearson correlations between the number of cell types per dataset, the number of cells per dataset, the median number of genes detected per cell, or the median number of reads per cell and wall time, **c**, or peak memory usage, **d**. Statistically significant correlations are shown in blue. Augur runtime scales approximately linearly with the number of cell types, while RAM usage scales with both the number of cell types and the number of cells. **e**, Scatterplots showing the relationships between wall time (top) and peak memory usage (bottom), and the dataset properties shown in **c-d**, for each of 22 scRNA-seq datasets in the compendium.

**Supplementary Figure 13.**
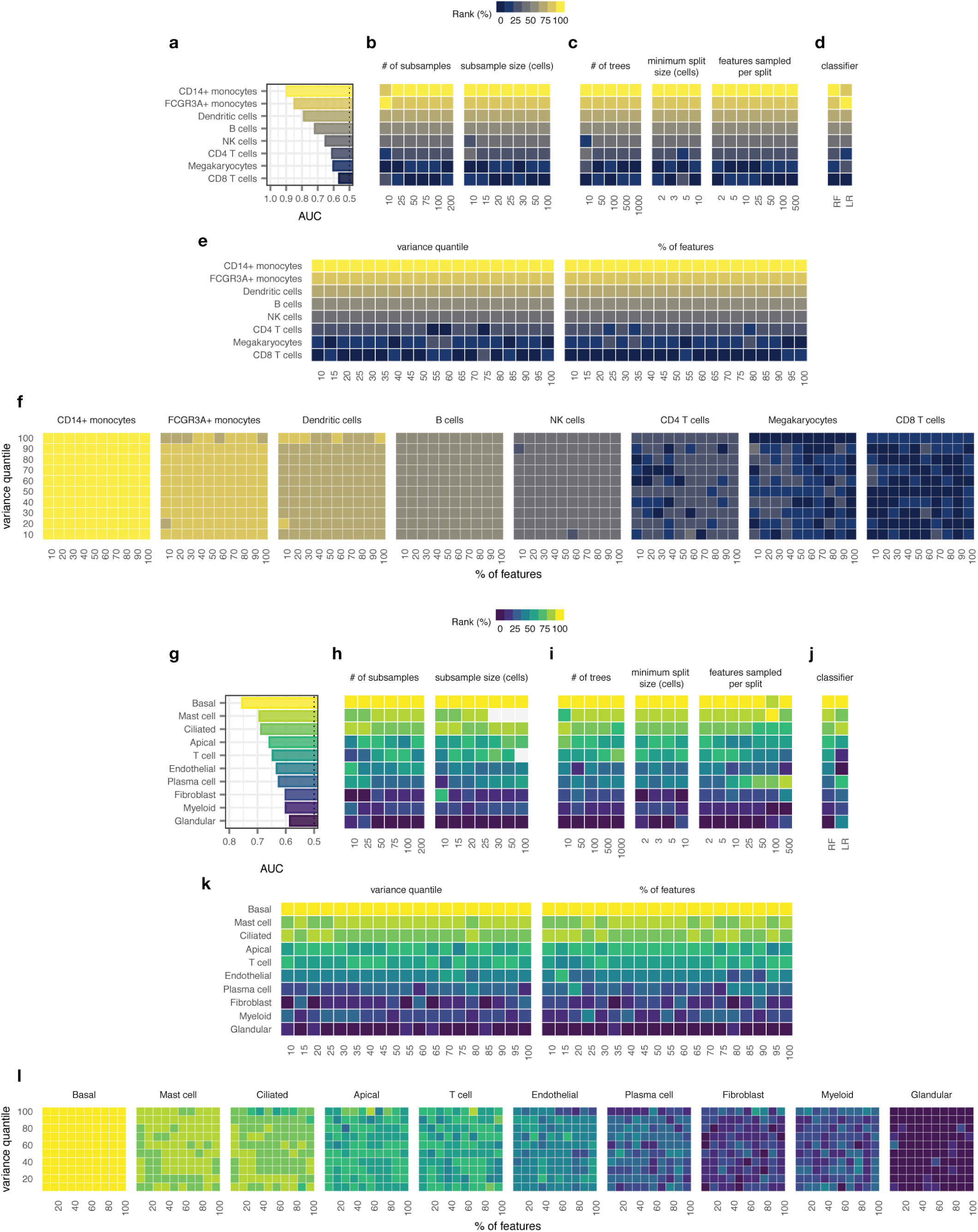
Robustness of cell type prioritization to Augur hyperparameters. **a-f**, Robustness of cell type prioritization for peripheral blood mononuclear cells stimulated with recombinant interferon-*β* in the Kang et al., 2018 dataset^4^. **g-l**, Robustness of cell type prioritization for ethmoid sinus cells from patients with chronic rhinosinusitis (CRS) in the Ordovas-Montanes et al., 2018 dataset^66^. **a** and **g**, Cell type prioritization (AUCs) under default parameters. **b** and **h**, Impact of varying the number of subsamples drawn from each cell type-specific gene expression matrix, or the size of each subsample, on cell type prioritization. **c** and **i**, Impact of varying random forest-specific hyperparameters, including the number of trees, minimum split size, and features sampled per split, on cell type prioritization. Grey tiles indicate sample sizes larger than the number of cells of that type in the dataset. **d** and **j**, Impact of replacing the random forest classifier (RF) with a L1-penalized logistic regression model (LR) on cell type prioritization. **e** and **k**, Impact of varying the proportion of highly variable genes selected from the original gene expression matrix, left, or the proportion of genes randomly selected in each subsample after the application of the variable gene filter, right, on cell type prioritization. **f** and **l**, Impact of simultaneously varying both the variable gene and random selection filters on cell type prioritization.

**Supplementary Figure 14.**
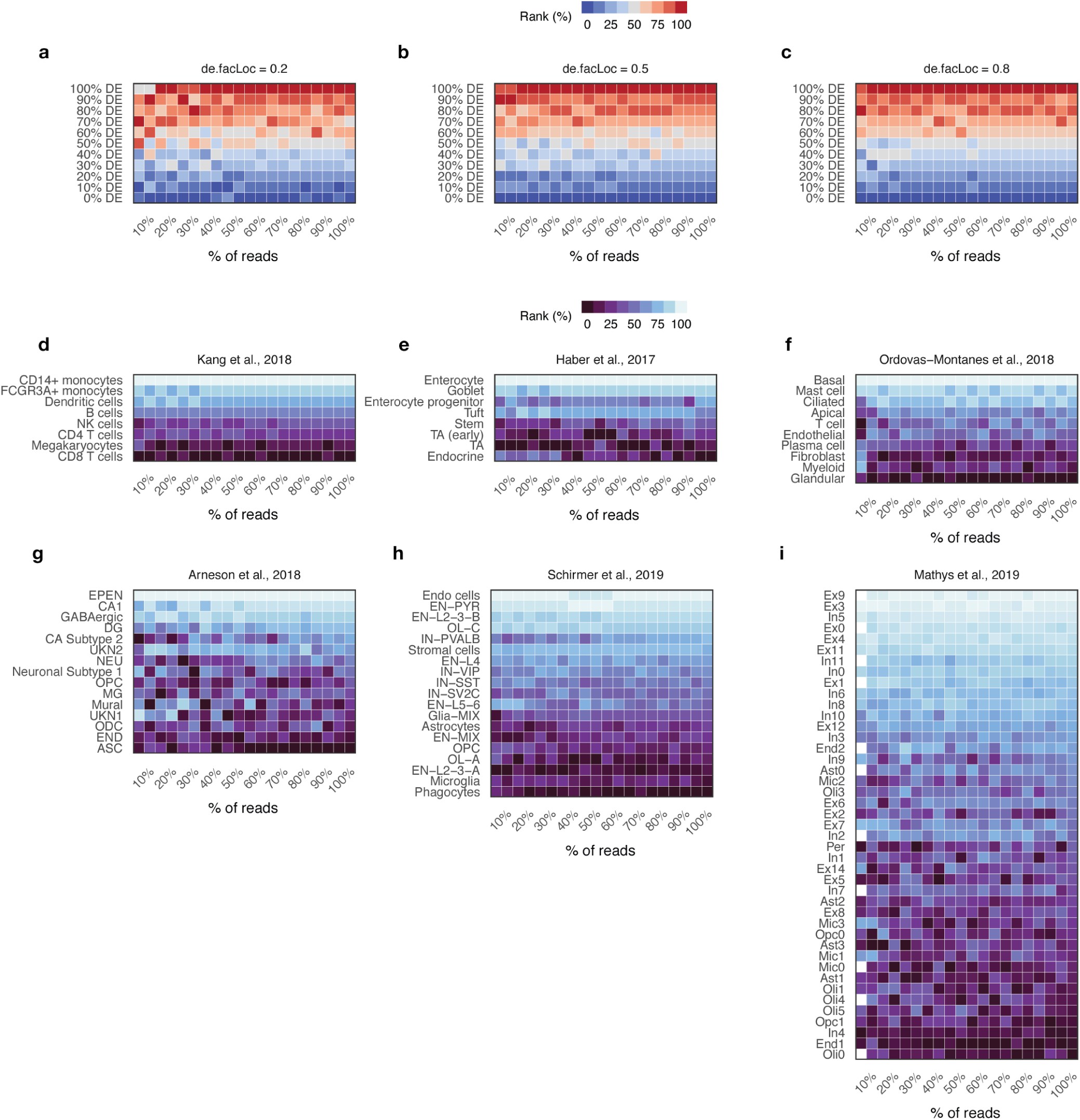
Impact of sequencing read depth on cell type prioritization. **a-c**, Augur cell type prioritization in simulated populations of *n* = 200 cells with between 0% and 100% of genes differentially expressed between experimental conditions, and read counts randomly downsampled from 100% to 5% of the original depth in 5% increments. Cell type prioritizations are robust to downsampling sequencing depth in simulated data; however, adequate depth becomes increasingly important with more subtle perturbations, shown by varying the location parameter of the differential expression factor log-normal distribution between 0.2, **a**; 0.5, **b**; and 0.8, **c**. **d-i**, Augur cell type prioritizations in six published scRNA-seq datasets with varying numbers of cell types, with read counts randomly downsampled from 100% to 5% of the original depth in 5% increments.

**Supplementary Figure 15.**
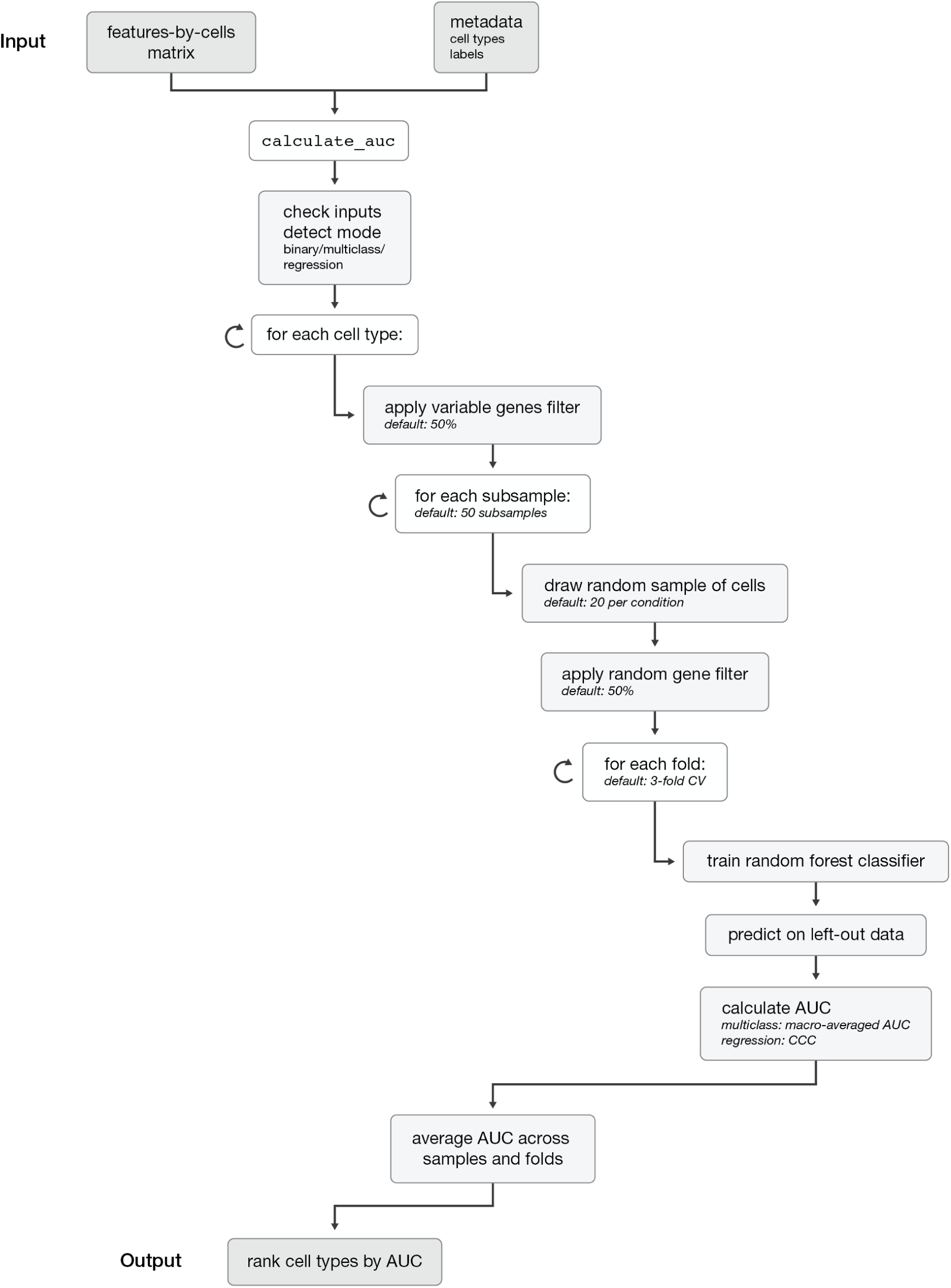
Overview of the Augur algorithm.

