## Supplementary Software for "Cell type prioritization in single-cell data": calculate_auc.R

```
#' Prioritize cell types involved in a biological process
#'
#' Prioritize cell types involved in a complex biological process by training a
#' machine-learning model to predict sample labels (e.g., disease vs. control,
#' treated vs. untreated, or time post-stimulus), and evaluate the performance 
#' of the model in cross-validation.
#'
#' If a \code{Seurat} object is provided as input, Augur will use the default
#' assay (i.e., whatever \link[Seurat]{GetAssayData} returns) as input. To
#' use a different assay, provide the expression matrix and metadata as input
#' separately, using the \code{input} and \code{meta} arguments.
#'
#' @param input a matrix, data frame, or \code{Seurat} or \code{monocle} object
#'   containing gene expression values (genes in rows, cells in columns) and,
#'   optionally, metadata about each cell
#' @param meta a data frame containing metadata about the \code{input}
#'   gene-by-cell matrix, at minimum containing the cell type for each cell
#'   and the labels (e.g., group, disease, timepoint); can be left as
#'   \code{NULL} if \code{input} is a \code{Seurat} or \code{monocle} object
#' @param label_col the column of the \code{meta} data frame, or the
#'   metadata container in the \code{Seurat} or \code{monocle} object, that
#'   contains condition labels (e.g., disease, timepoint) for each cell in the
#'   gene-by-cell expression matrix; defaults to \code{label}
#' @param cell_type_col the column of the \code{meta} data frame, or the
#'   metadata container in the \code{Seurat}/\code{monocle} object, that
#'   contains cell type labels for each cell in the gene-by-cell expression
#'   matrix; defaults to \code{cell_type}
#' @param n_subsamples the number of random subsamples of fixed size to
#'   draw from the complete dataset, for each cell type; defaults to \code{50}.
#'   Set to \code{0} to omit subsampling altogether,
#'   calculating performance on the entire dataset, but note that this may
#'   introduce bias due to cell type or label class imbalance
#' @param subsample_size the number of cells per type to subsample randomly from
#'   each experimental condition, if \code{n_subsamples} is greater than 1; 
#'   defaults to \code{20}
#' @param folds the number of folds of cross-validation to run; defaults to
#'   \code{3}. Be careful changing this parameter without also changing
#'   \code{subsample_size}
#' @param min_cells the minimum number of cells for a particular cell type in
#'   each condition in order to retain that type for analysis;
#'   defaults to \code{subsample_size}
#' @param var_quantile the quantile of highly variable genes to retain for
#'   each cell type using the variable gene filter (\link{select_variance});
#'   defaults to \code{0.5}
#' @param feature_perc the proportion of genes that are randomly selected as
#'   features for input to the classifier in each subsample using the 
#'   random gene filter (\link{select_random}); defaults to \code{0.5}
#' @param n_threads the number of threads to use for parallelization;
#'   defaults to \code{4}.
#' @param show_progress if \code{TRUE}, display a progress bar for the analysis
#'   with estimated time remaining
#' @param classifier the classifier to use in calculating area under the curve,
#'   one of \code{"rf"} (random forest) or \code{"lr"} (logistic regression);
#'   defaults to \code{"rf"}, which is the recommended setting
#' @param rf_params for \code{classifier} == \code{"rf"}, a list of parameters
#'   for the random forest models, containing the following items (see 
#'   \link[parsnip]{rand_forest} from the \code{parsnip} package): 
#'   \describe{
#'     \item{"mtry"}{the number of features randomly sampled at each split 
#'       in the random forest classifier; defaults to \code{2}}
#'     \item{"trees"}{the number of trees in the random forest classifier;
#'       defaults to \code{100}}
#'     \item{"min_n"}{the minimum number of observations to split a node in the
#'       random forest classifier; defaults to \code{NULL}}
#'     \item{"importance"}{the method of calculating feature importances
#'       to use; defaults to \code{"accuracy"}; can also specify \code{"gini"}}
#'   }
#' @param lr_params for \code{classifier} == \code{"lr"}, a list of parameters
#'   for the logistic regression models, containing the following items (see 
#'   \link[parsnip]{logistic_reg} from the \code{parsnip} package):  
#'   \describe{
#'     \item{"mixture"}{the proportion of L1 regularization in the model;
#'       defaults to \code{1}}
#'     \item{"penalty"}{the total amount of regularization in the model; 
#'       defaults to \code{"auto"}, which uses \link[glmnet]{cv.glmnet} to set 
#'       the penalty}
#'   }
#' 
#' @return a list of class \code{"Augur"}, containing the following items:
#' \enumerate{
#'   \item \code{X}: the numeric matrix (or data frame or sparse matrix, 
#'     depending on the input) containing gene expression values for each cell 
#'     in the dataset
#'   \item \code{y}: the vector of experimental condition labels being predicted
#'   \item \code{cell_types}: the vector of cell type labels
#'   \item \code{parameters}: the parameters provided to this function as input
#'   \item \code{results}: the area under the curve for each cell type, in each
#'     fold, in each subsample, in the comparison of interest, as well as a
#'     series of other classification metrics
#'   \item \code{feature_importance}: the importance of each feature for
#'     calculating the AUC, above. For random forest classifiers, this is the
#'     mean decrease in accuracy or Gini index. For logistic regression 
#'     classifiers, this is the standardized regression coefficients, computed
#'     using the Agresti method
#'   \item \code{AUC}: a summary of the mean AUC for each cell type (for 
#'     continuous experimental conditions, this is replaced by a \code{CCC} 
#'     item that records the mean concordance correlation coefficient for each 
#'     cell type)
#' }
#'
#' @importFrom dplyr do sample_n group_by ungroup tibble mutate select bind_rows
#'   pull rename n_distinct arrange desc filter summarise
#' @importFrom tibble repair_names rownames_to_column
#' @importFrom purrr map map2 map_lgl pmap map2_df
#' @importFrom magrittr %>% %<>% extract extract2 set_rownames set_colnames
#' @importFrom parsnip set_engine logistic_reg rand_forest fit translate
#' @importFrom rsample assessment analysis
#' @importFrom recipes prepper bake recipe
#' @importFrom yardstick metric_set accuracy precision recall sens spec npv
#'   ppv roc_auc ccc huber_loss_pseudo huber_loss mae mape mase
#'   rpd rpiq rsq_trad rsq smape rmse
#' @importFrom stats setNames predict sd
#' @importFrom methods is
#' @importFrom sparseMatrixStats colVars
#' @importFrom pbmcapply pbmclapply
#' @importFrom parallel mclapply
#' @importFrom tester is_numeric_matrix is_numeric_dataframe
#' @import Matrix
#'
#' @export
calculate_auc = function(input,
                         meta = NULL,
                         label_col = "label",
                         cell_type_col = "cell_type",
                         n_subsamples = 50,
                         subsample_size = 20,
                         folds = 3,
                         min_cells = NULL,
                         var_quantile = 0.5,
                         feature_perc = 0.5,
                         n_threads = 4,
                         show_progress = T,
                         classifier = c("rf", "lr"),
                         # random forest parameters
                         rf_params = list(trees = 100,
                                          mtry = 2,
                                          min_n = NULL,
                                          importance = 'accuracy'),
                         # logistic regression parameters
                         lr_params = list(mixture = 1, penalty = 'auto')
) {
  # check arguments
  classifier = match.arg(classifier)
  # check number of folds/subsample size are compatible (n > 1 in every fold)
  if (n_subsamples > 1 & subsample_size / folds < 2) {
    stop("subsample_size / n_folds must be greater than or equal to 2")
  }
  # make sure regression is only used with random rand_forest
  if (is.null(min_cells)) {
    min_cells = subsample_size
  }

  # check glmnet is installed for logistic regression
  if (!requireNamespace("glmnet", quietly = TRUE)) {
    stop("install \"glmnet\" R package to run Augur with logistic regression ",
         "classifier", call. = FALSE)
  }
  
  # extract cell types and label from metadata
  if ("Seurat" %in% class(input)) {
    # confirm Seurat is installed
    if (!requireNamespace("Seurat", quietly = TRUE)) {
      stop("install \"Seurat\" R package for Augur compatibility with ",
           "input Seurat object", call. = FALSE)
    }

    meta = input@meta.data %>%
      droplevels()
    cell_types = meta[[cell_type_col]]
    labels = meta[[label_col]]
    expr = Seurat::GetAssayData(input)
  } else if ("cell_data_set" %in% class(input)) {
    # confirm monocle3 is installed
    if (!requireNamespace("monocle3", quietly = TRUE)) {
      stop("install \"monocle3\" R package for Augur compatibility with ",
           "input monocle3 object", call. = FALSE)
    }

    meta = monocle3::pData(input) %>%
      droplevels() %>%
      as.data.frame()
    cell_types = meta[[cell_type_col]]
    labels = meta[[label_col]]
    expr = monocle3::exprs(input)
  } else if ("SingleCellExperiment" %in% class(input)){ 
    # confirm SingleCellExperiment is installed
    if (!requireNamespace("SingleCellExperiment", quietly = TRUE)) {
      stop("install \"SingleCellExperiment\" R package for Augur ",
           "compatibility with input SingleCellExperiment object", 
           call. = FALSE)
    }
    
    meta = SummarizedExperiment::colData(input) %>%
      droplevels() %>%
      as.data.frame()
    cell_types = meta[[cell_type_col]]
    labels = meta[[label_col]]
    expr = SummarizedExperiment::assay(input)
  } else {
    # data frame or matrix plus metadata data frame
    if (is.null(meta)) {
      stop("must provide metadata if not supplying a Seurat or monocle object")
    }

    # check if input is sparse matrix or numberic matrix/df
    valid_input = is(input, 'sparseMatrix') ||
      is_numeric_matrix(input) ||
      is_numeric_dataframe(input)
    if (!valid_input)
      stop("input must be Seurat, monocle, sparse matrix, numeric matrix, or ",
           "numeric data frame")

    # extract columns
    expr = input
    meta %<>% droplevels()
    cell_types = meta[[cell_type_col]]
    labels = meta[[label_col]]
  } 

  # check dimensions are non-zero
  if (length(dim(expr)) != 2 || !all(dim(expr) > 0)) {
    stop("expression matrix has at least one dimension of size zero")
  }

  # check dimensions match
  n_cells1 = nrow(meta)
  n_cells2 = ncol(expr)
  if (n_cells1 != n_cells2) {
    stop("number of cells in metadata (", n_cells1, ") does not match number ",
         "of cells in expression (", n_cells2, ")")
  }

  # check at least two labels
  if (n_distinct(labels) == 1) {
    stop("only one label provided: ", unique(labels))
  }

  # check for missing labels or cell types
  if (any(is.na(labels))) {
    stop("labels contain ", sum(is.na(labels)), "missing values")
  }
  if (any(is.na(cell_types))) {
    stop("cell types contain ", sum(is.na(cell_types)), "missing values")
  }

  # make sure `label` is not a rowname in `input` (feature in RF)
  if ("label" %in% rownames(expr)) {
    warning("row `label` exists in input; changing ...")
    to_fix = which(rownames(expr) == "label")
    rownames(expr)[to_fix] = paste0("label", seq_along(rownames(expr)[to_fix]))
  }

  # fix column names that ranger can't handle
  ## (here, they are still rows)
  rf_engine = "randomForest" ## set to randomForest - don't use ranger
  if (classifier == "rf" && rf_engine == "ranger") {
    invalid_rows = any(grepl("-|\\.|\\(|\\)", rownames(expr)))
    if (invalid_rows) {
      warning("classifier `rf` with engine `ranger` cannot handle characters ",
              "-.() in column names; replacing ...")
      expr %<>% set_rownames(gsub("-|\\.|\\(|\\)", "", rownames(.)))
    }
  }

  # remove missing values
  missing = is.na(expr)
  if (any(missing)) {
    stop("matrix contains ", sum(missing), "missing values")
  }

  # detect mode
  if (is.numeric(labels)) {
    mode = "regression"
    multiclass = F

    # throw a warning if there are only three or less values
    if (n_distinct(labels) <= 3) {
      warning("doing regression with only ", n_distinct(labels),
              " unique values")
    }
  } else {
    mode = "classification"

    # check whether we are working with multiclass data
    multiclass = n_distinct(labels) > 2
    # check classifier is compatible with classification problem
    if (multiclass & classifier == "lr") {
      stop("multi-class classification with classifier = 'lr' is currently not ",
           "supported in tidymodels `logistic_reg`")
    }

    # make sure y is a factor if doing classification
    if (!is.factor(labels)) {
      warning("coercing labels to factor ...")
      labels %<>% as.factor()
    }
  }

  # check if showing progress or not
  if (show_progress == T) {
    apply_fun = pbmclapply
  } else {
    apply_fun = mclapply
  }

  # iterate over cell type clusters
  res = apply_fun(unique(cell_types),
    mc.cores = n_threads, function(cell_type) {
      # skip this cell type if there aren't enough cells
      y = labels[cell_types == cell_type]
      if (mode == 'classification') {
        if (min(table(y)) < min_cells) {
          warning("skipping cell type ", cell_type,
                  ": minimum number of cells (", min(table(y)),
                  ") is less than ", min_cells)
          return(list())
        }
      } else if (mode == 'regression') {
        if (length(y) < min_cells) {
          warning("skipping cell type ", cell_type,
                  ": total number of cells (", length(y),
                  ") is less than ", min_cells)
          return(list())
        }
      }
      # subset the entire expression matrix to this cell type
      X = expr[, cell_types == cell_type]

      # select features by variance
      min_features_for_selection = 1000
      if (nrow(X) >= min_features_for_selection) {
        X %<>% select_variance(var_quantile, filter_negative_residuals = F)
      }

      # set up subsamples and results bin
      tmp_results = data.frame()
      tmp_importances = data.frame()

      n_iter = ifelse(n_subsamples < 1, 1, n_subsamples)
      for (subsample_idx in seq_len(n_iter)) {
        # seed RNG for reproducibility
        set.seed(subsample_idx)

        # optionally, skip the subsampling process
        if (n_subsamples < 1) {
          # randomly select features
          if (nrow(X) >= min_features_for_selection &
              feature_perc < 1) {
            X0 = select_random(X, feature_perc)
          } else {
            X0 = X
          }
          X0 %<>%
            t() %>%
            as.matrix() %>%
            as.data.frame() %>%
            # fix any non-unique columns
            repair_names() %>%
            # add labels
            mutate(label = y)
        } else {
          if (mode == 'regression') {
            subsample_idxs = data.frame(label = y,
                                        position = seq_along(y)) %>%
              do(sample_n(., subsample_size)) %>%
              pull(position)
          } else {
            subsample_idxs = data.frame(label = y,
                                        position = seq_along(y)) %>%
              group_by(label) %>%
              do(sample_n(., subsample_size)) %>%
              pull(position)
          }
          y0 = y[subsample_idxs]
          # randomly select features
          if (nrow(X) >= min_features_for_selection &
              feature_perc < 1) {
            X0 = select_random(X, feature_perc)
          } else {
            X0 = X
          }
          # coerce to data frame
          X0 %<>%
            extract(, subsample_idxs) %>%
            t() %>%
            extract(, colVars(.) > 0) %>%
            as.matrix() %>%
            as.data.frame() %>%
            # fix any non-unique columns
            repair_names() %>%
            # convert labels back to a factor
            mutate(label = y0)
        }

        # set up model
        if (classifier == "rf") {
          importance = T
          if (rf_engine == "ranger")
            importance = "impurity"
          clf = rand_forest(trees = !!rf_params$trees,
                            mtry = !!rf_params$mtry,
                            min_n = !!rf_params$min_n,
                            mode = mode) %>%
            set_engine(rf_engine, seed = 1, importance = T, localImp = T)
        } else if (classifier == "lr") {
          family = ifelse(multiclass, 'multinomial', 'binomial')

          if (is.null(lr_penalty) || lr_penalty == 'auto') {
            # first, get optimized penalty for dataset
            lr_penalty = withCallingHandlers({
              glmnet::cv.glmnet(X0 %>%
                          ungroup() %>%
                          select(-label) %>%
                          as.matrix() %>%
                          extract(, setdiff(colnames(.), 'label')),
                        X0$label,
                        nfolds = folds,
                        family = family) %>%
                extract2("lambda.1se")
            }, warning = function(w) {
              if (grepl("dangerous ground", conditionMessage(w)))
                invokeRestart("muffleWarning")
            })
          }

          clf = logistic_reg(mixture = lr_mixture,
                             penalty = lr_penalty,
                             mode = 'classification') %>%
            set_engine('glmnet', family = family)
        } else {
          stop("invalid classifier: ", classifier)
        }

        # fit models in cross-validation
        if (mode == "classification") {
          cv = vfold_cv(X0, v = folds, strata = 'label')
        } else {
          cv = vfold_cv(X0, v = folds)
        }
        withCallingHandlers({
          folded = cv %>%
            mutate(
              recipes = splits %>%
                map(~ prepper(., recipe = recipe(.$data, label ~ .))),
              test_data = splits %>% map(analysis),
              fits = map2(
                recipes,
                test_data,
                ~ fit(
                  clf,
                  label ~ .,
                  data = bake(object = .x, new_data = .y)
                )
              )
            )
        }, warning = function(w) {
          if (grepl("dangerous ground", conditionMessage(w)))
            invokeRestart("muffleWarning")
        })

        # predict on the left-out data
        retrieve_class_preds = function(split, recipe, model) {
          test = bake(recipe, assessment(split))
          tbl = tibble(
            true = test$label,
            pred = predict(model, test)$.pred_class,
            prob = predict(model, test, type = 'prob')) %>%
            # convert prob from nested df to columns
            cbind(.$prob) %>%
            select(-prob)
          return(tbl)
        }
        retrieve_reg_preds = function(split, recipe, model) {
          test = bake(recipe, assessment(split))
          tbl = tibble(
            true = test$label,
            pred = predict(model, test)$.pred)
          return(tbl)
        }
        predictions = folded %>%
          mutate(
            pred = list(
              splits,
              recipes,
              fits
            )
          )
        if (mode == 'regression') {
          predictions = predictions %>%
            mutate(pred = pmap(pred, retrieve_reg_preds))
        } else {
          predictions = predictions %>%
            mutate(pred = pmap(pred, retrieve_class_preds))
        }

        # evalulate the predictions
        if (mode == 'regression') {
          multi_metric = metric_set(ccc, huber_loss_pseudo, huber_loss,
            mae, mape, mase, rpd, rpiq, rsq_trad, rsq, smape, rmse)
        } else {
          roc_auc_custom = function(...) roc_auc(...,
                                                 options = list(direction = "<"))
          class(roc_auc_custom) = class(roc_auc)
          multi_metric = metric_set(accuracy, precision, recall, sens,
                                    spec, npv, ppv,
                                    roc_auc_custom)
        }

        prob_select = 3
        if (mode == "classification") {
          estimator = ifelse(multiclass, "macro", "binary")
          if (multiclass)
            prob_select = seq(3, 3 + n_distinct(labels) - 1)
          metric_fun = function(x)
            multi_metric(x,
                         truth = true,
                         estimate = pred,
                         prob_select,
                         estimator = estimator)
        } else {
          metric_fun = function(x)
            multi_metric(x,
                         truth = true,
                         estimate = pred,
                         prob_select)
        }
        eval = predictions %>%
          mutate(
            metrics = pred %>%
              map(metric_fun)
          ) %>%
          extract2('metrics')

        # clean up the results
        result = eval %>%
          map2_df(., names(.), ~ mutate(.x, fold = .y)) %>%
          set_colnames(gsub("\\.", "", colnames(.))) %>%
          mutate(cell_type = cell_type,
                 subsample_idx = subsample_idx)

        # also calculate feature importance
        importance = NULL
        if (classifier == "rf") {
          importance_name = 'importance'
          if (mode == 'regression') {
            if (rf_params$importance == 'accuracy') {
              impval_name = "%IncMSE"
            } else {
              impval_name = 'IncNodePurity'
            }
          } else {
            if (rf_params$importance == 'accuracy') {
              impval_name = 'MeanDecreaseAccuracy'
            } else {
              impval_name = 'MeanDecreaseGini'
            }
          }
          if (rf_engine == "ranger") {
            importance_name = "variable.importance"
            impval_name == ".x[[i]]"
          }

          importance = folded %>%
            pull(fits) %>%
            map("fit") %>%
            map(importance_name) %>%
            map(as.data.frame) %>%
            map(~ rownames_to_column(., 'gene')) %>%
            map2_df(names(.), ~ mutate(.x, fold = .y)) %>%
            mutate(cell_type = cell_type,
                   subsample_idx = subsample_idx) %>%
            dplyr::rename(importance = impval_name) %>%
            # rearrange columns
            dplyr::select(cell_type, subsample_idx, fold, gene, importance)
        } else if (classifier == "lr") {
          # standardized coefficients with Agresti method
          # cf. https://think-lab.github.io/d/205/#3
          coefs = folded %>%
            pull(fits) %>%
            map("fit") %>%
            map(~ as.matrix(coef(., s = lr_penalty)))
          sds = folded %>%
            pull(splits) %>%
            map('data') %>%
            # omit labels
            map(~ extract(., , -ncol(.))) %>%
            map(~ apply(., 2, sd))
          std_coefs = map(seq_len(folds), ~ coefs[[.]][-1, 1] * sds[[.]])
          importance = std_coefs %>%
            map(~ data.frame(gene = names(.), std_coef = .)) %>%
            setNames(seq_len(folds)) %>%
            map2_df(names(.), ~ mutate(.x, fold = .y)) %>%
            mutate(cell_type = cell_type,
                   subsample_idx = subsample_idx) %>%
            # rearrange columns
            dplyr::select(cell_type, subsample_idx, fold, gene, std_coef)
        }

        # rearrange columns
        result %<>%
          dplyr::select(cell_type, subsample_idx, fold, metric, estimator,
                        estimate)

        # add to results
        tmp_results %<>% bind_rows(result)
        tmp_importances %<>% bind_rows(importance)
      }
      list(results = tmp_results, importances = tmp_importances)
    }
  )

  # ignore warnings from yardstick
  if (any(map_lgl(res, ~ "warning" %in% class(.)))) {
    res = res$value
  }

  # make sure at least one cell type worked
  if (all(lengths(res) == 0))
    stop("no cell type had at least ", min_cells, " cells in all conditions")

  # summarise AUCs (or CCCs) per cell type
  if (mode == "classification") {
    AUCs = res %>%
      map("results") %>%
      bind_rows() %>%
      filter(metric == "roc_auc") %>%
      group_by(cell_type, subsample_idx) %>%
      summarise(estimate = mean(estimate)) %>%
      ungroup() %>%
      group_by(cell_type) %>%
      summarise(auc = mean(estimate)) %>%
      ungroup() %>%
      arrange(desc(auc))
  } else if (mode == "regression") {
    CCCs = res %>%
      map("results") %>%
      bind_rows() %>%
      filter(metric == "ccc") %>%
      group_by(cell_type, subsample_idx) %>%
      summarise(estimate = mean(estimate)) %>%
      ungroup() %>%
      group_by(cell_type) %>%
      summarise(ccc = mean(estimate)) %>%
      ungroup() %>%
      arrange(desc(ccc))
  }

  # clean up results across lapply folds
  feature_importances = res %>%
    map("importances") %>%
    bind_rows()
  results = res %>%
    map("results") %>%
    bind_rows()

  # create Augur object
  params = list(
    n_subsamples = n_subsamples,
    subsample_size = subsample_size,
    folds = folds,
    min_cells = min_cells,
    var_quantile = var_quantile,
    feature_perc = feature_perc,
    n_threads = n_threads,
    classifier = classifier
  )
  if (classifier == "rf")
    params$rf_params = rf_params
  if (classifier == "lr")
    params$lr_params = lr_params
  obj = list(
    X = expr,
    y = labels,
    cell_types = cell_types,
    parameters = params,
    results = results,
    feature_importance = feature_importances
  )
  if (mode == "classification") {
    obj$AUC = AUCs
  } else if (mode == "regression") {
    obj$CCC = CCCs
  }
  
  return(obj)
}
```
