## Supplementary Software for "Cell type prioritization in single-cell data": custom_vfold_cv.R

```
# v-fold cross-validation
# (copied from rsample package, with edits for >7-class classification)
 
#' @import rsample
#' @importFrom tidyselect vars_select
#' @importFrom rlang enquo
vfold_cv <- function(data, v = 10, repeats = 1, strata = NULL, breaks = 4,
                     ...) {
  
  if(!missing(strata)) {
    strata <- vars_select(names(data), !!enquo(strata))
    if(length(strata) == 0) strata <- NULL
  }
  
  strata_check(strata, names(data))
  
  if (repeats == 1) {
    split_objs <- vfold_splits(data = data, v = v, strata = strata, 
                               breaks = breaks)
  } else {
    for (i in 1:repeats) {
      tmp <- vfold_splits(data = data, v = v, strata = strata)
      tmp$id2 <- tmp$id
      tmp$id <- names0(repeats, "Repeat")[i]
      split_objs <- if (i == 1)
        tmp
      else
        rbind(split_objs, tmp)
    }
  }
  
  ## We remove the holdout indicies since it will save space and we can
  ## derive them later when they are needed.
  
  split_objs$splits <- map(split_objs$splits, rm_out)
  
  ## Save some overall information
  
  cv_att <- list(v = v, repeats = repeats, strata = !is.null(strata))
  
  new_rset(splits = split_objs$splits,
           ids = split_objs[, grepl("^id", names(split_objs))],
           attrib = cv_att,
           subclass = c("vfold_cv", "rset"))
}

# Get the indices of the analysis set from the assessment set
vfold_complement <- function(ind, n) {
  list(analysis = setdiff(1:n, ind),
       assessment = ind)
}

#' @import rsample
#' @importFrom tibble tibble
#' @importFrom purrr map
#' @importFrom dplyr bind_rows
vfold_splits <- function(data, v = 10, strata = NULL, breaks = 4) {
  if (!is.numeric(v) || length(v) != 1)
    stop("`v` must be a single integer.", call. = FALSE)
  
  n <- nrow(data)
  if (is.null(strata)) {
    folds <- sample(rep(1:v, length.out = n))
    idx <- seq_len(n)
    indices <- split(idx, folds)
  } else {
    stratas <- tibble::tibble(idx = 1:n,
                              strata = make_strata(getElement(data, strata),
                                                   breaks = breaks,
                                                   pool = 0.1))
    stratas <- split(stratas, stratas$strata)
    stratas <- purrr::map(stratas, add_vfolds, v = v)
    stratas <- dplyr::bind_rows(stratas)
    indices <- split(stratas$idx, stratas$folds)
  }
  
  indices <- lapply(indices, vfold_complement, n = n)
  
  split_objs <- purrr::map(indices, make_splits, data = data, 
                           class = "vfold_split")
  tibble::tibble(splits = split_objs,
                 id = names0(length(split_objs), "Fold"))
}

add_vfolds <- function(x, v) {
  x$folds <- sample(rep(1:v, length.out = nrow(x)))
  x
}

strata_check <- function(strata, vars) {
  if (!is.null(strata)) {
    if (!is.character(strata) | length(strata) != 1)
      stop("`strata` should be a single character value", call. = FALSE)
    if (!(strata %in% vars))
      stop(strata, " is not in `data`")
  }  
  invisible(NULL)
}

make_splits <- function(ind, data, class = NULL) {
  res <- rsplit(data, ind$analysis,  ind$assessment)
  if (!is.null(class))
    res <- add_class(res, class)
  res
}

rsplit <- function(data, in_id, out_id) {
  if (!is.data.frame(data) & !is.matrix(data))
    stop("`data` must be a data frame.", call. = FALSE)
  
  if (!is.integer(in_id) | any(in_id < 1))
    stop("`in_id` must be a positive integer vector.", call. = FALSE)
  
  if(!all(is.na(out_id))) {
    if (!is.integer(out_id) | any(out_id < 1))
      stop("`out_id` must be a positive integer vector.", call. = FALSE)
  }
  
  if (length(in_id) == 0)
    stop("At least one row should be selected for the analysis set.",
         call. = FALSE)
  
  structure(
    list(
      data = data,
      in_id = in_id,
      out_id = out_id
    ),
    class = "rsplit"
  )
}

add_class <- function(x, cls, at_end = TRUE) {
  class(x) <- if (at_end)
    c(class(x), cls)
  else
    c(cls, class(x))
  x
}

names0 <- function (num, prefix = "x") {
  if (num < 1) 
    stop("`num` should be > 0", call. = FALSE)
  ind <- format(1:num)
  ind <- gsub(" ", "0", ind)
  paste0(prefix, ind)
}

## This will remove the assessment indices from an rsplit object
rm_out <- function(x) {
  x$out_id <- NA
  x
}

#' @importFrom tibble is_tibble as_tibble tibble
#' @importFrom dplyr bind_cols
# `splits`` should be either a list or a tibble with a single column
#  called "splits"
# `ids`` should be either a character vector or a tibble with 
#  one or more columns that begin with "id" 
new_rset <-  function(splits, ids, attrib = NULL, 
                      subclass = character()) {
  stopifnot(is.list(splits))
  if (!is_tibble(ids)) {
    ids <- tibble(id = ids)
  } else {
    if (!all(grepl("^id", names(ids))))
      stop("The `ids` tibble column names should start with 'id'",
           call. = FALSE)
  }
  either_type <- function(x)
    is.character(x) | is.factor(x)
  ch_check <- vapply(ids, either_type, c(logical = TRUE))
  if(!all(ch_check))
    stop("All ID columns should be character or factor ",
         "vectors.", call. = FALSE)
  
  if (!is_tibble(splits)) {
    splits <- tibble(splits = splits)
  } else {
    if(ncol(splits) > 1 | names(splits)[1] != "splits")
      stop("The `splits` tibble should have a single column ", 
           "named `splits`.", call. = FALSE)
  }
  
  if (nrow(ids) != nrow(splits))
    stop("Split and ID vectors have different lengths.",
         call. = FALSE)  
  
  # Create another element to the splits that is a tibble containing
  # an identifer for each id column so that, in isolation, the resample
  # id can be known just based on the `rsplit` object. This can then be
  # accessed using the `labels` methof for `rsplits`
  
  splits$splits <- map2(splits$splits, split(ids, 1:nrow(ids)), add_id)
  
  res <- bind_cols(splits, ids)
  
  if (!is.null(attrib)) {
    if (any(names(attrib) == ""))
      stop("`attrib` should be a fully named list.",
           call. = FALSE)
    for (i in names(attrib))
      attr(res, i) <- attrib[[i]]
  }
  
  if (length(subclass) > 0)
    res <- add_class(res, cls = subclass, at_end = FALSE)
  
  res
}


add_id <- function(split, id) {
  split$id <- id
  split
}
```
