## Supplementary Software for "Cell type prioritization in single-cell data": sc_sim.R

```
#' A simulated single-cell RNA-seq dataset
#' 
#' A toy simulated scRNA-seq dataset, used to demonstrate Augur. 
#' The dataset contains three populations of cells (CellTypeA, CellTypeB, and
#' CellTypeC), each represented by 200 cells.
#' Each cell type has approximately half of its cells in one of two experimental
#' conditions, denoted as "treatment" and "control."
#' The cell types have varying proportions of genes differentially expressed in
#' response to the treatment: cell type A has 5% of its genes DE, 
#' cell type B has 25% of its genes DE, and cell type C has 50% of its genes
#' DE. 
#' 
#' The dataset was simulated with splatter, with simulation parameters estimated
#' from the Kang et al., 2018 dataset. 
#' The data is provided as a Seurat object. 
#' 
#' @docType data
#' @usage data(sc_sim)
#' @format a Seurat object with dimensions 15,697 genes x 600 cells
"sc_sim"
```
