## Supplementary Software for "Cell type prioritization in single-cell data": select_random.R

```
#' Random feature selection
#' 
#' Perform feature selection on a single-cell feature matrix (e.g., gene 
#' expression) by randomly removing a specified proportion of features.
#' 
#' @param mat a single-cell matrix to be filtered, with features (genes) in rows
#'   and cells in columns
#' @param feature_perc percentage of features (genes) chosen at random following
#'   the technical variance filter
#' 
#' @return the filtered matrix (or, if \code{feature_perc == 1}, the input 
#'   matrix)
#'
#' @importFrom magrittr %<>% extract
#' @export
select_random = function(mat, feature_perc = 0.5) {
  if (feature_perc < 1) {
    features = rownames(mat)
    keep = sample(features, floor(nrow(mat) * feature_perc))
    mat %<>% extract(keep, )
  }
  return(mat)
}
```
