## Supplementary Software for "Cell type prioritization in single-cell data": select_variance.R

```
#' Feature selection based on variance
#'
#' Perform feature selection on a single-cell feature matrix (e.g., gene
#' expression) by first removing constant features, then removing features with
#' lower than expected variance, as quantified by the residuals from a loess
#' regression of feature (gene) coefficient of variation against mean
#' expression.
#'
#' @param mat a single-cell matrix to be filtered, with features (genes) in rows
#'   and cells in columns
#' @param var_quantile the quantile below which features will be filtered,
#'   based on their residuals in a loess model; defaults to \code{0.5}
#' @param filter_negative_residuals if \code{TRUE}, filter residuals at a fixed
#'   threshold of zero, instead of \code{var_quantile}
#'
#' @return the filtered matrix (or, if \code{var_quantile == 1} and
#'   \code{filter_negative_residuals == FALSE}, the input matrix)
#'
#' @importFrom dplyr between
#' @importFrom Matrix rowMeans rowSums
#' @importFrom lmtest coxtest
#' @importFrom stats loess quantile
#' @importFrom sparseMatrixStats rowSds
#' @importFrom magrittr %<>% extract
#'
#' @export
select_variance = function(mat,
                           var_quantile = 0.5,
                           filter_negative_residuals = FALSE) {
  # first, remove features with constant variance
  sds = rowSds(mat)
  # constant variance can cause an NA using this implementation, fix
  sds[is.na(sds)] <- 0
  mat %<>% extract(sds > 0, )

  # next, if var_quantile < 1, fit loess model
  if (var_quantile < 1 | filter_negative_residuals) {
    # calculate mean and coefficient of variation
    means = rowMeans(mat)
    sds %<>% extract(. > 0)
    cvs = means / sds
    # clip outliers, get best prediction model, get residuals
    lower = quantile(cvs, 0.01)
    upper = quantile(cvs, 0.99)
    keep = between(cvs, lower, upper)
    cv0 = cvs[keep]
    mean0 = means[keep]
    if (any(mean0 < 0)) {
      # if there are negative values, don't bother comparing to log-transformed
      # means - fit on normalized data directly
      model = loess(cv0 ~ mean0)
    } else {
      # use a cox test to guess whether we should be fitting the raw or
      # log-transformed means
      fit1 = loess(cv0 ~ mean0)
      fit2 = loess(cv0 ~ log(mean0))
      cox = coxtest(fit1, fit2)
      probs = cox$`Pr(>|z|)`
      if (probs[1] < probs[2]) {
        model = fit1
      } else {
        model = fit2
      }
    }
    genes = rownames(mat)[keep]
    residuals = setNames(model$residuals, genes)

    # select features by quantile (or override w/positive residuals)
    if (filter_negative_residuals == T) {
      genes = names(residuals)[residuals > 0]
    } else {
      genes = names(residuals)[residuals > quantile(residuals, var_quantile)]
    }
    mat %<>% extract(genes, )
  }

  return(mat)
}
```
