## Supplementary Software for "Cell type prioritization in single-cell data": README.md

# README

Augur is an R package to prioritize cell types involved in the response to an experimental perturbation within high-dimensional single-cell data. 
The intuition underlying Augur is that cells undergoing a profound response to a given experimental stimulus become more separable, in the space of molecular measurements, than cells that remain unaffected by the stimulus. 
Augur quantifies this separability by asking how readily the experimental sample labels associated with each cell (e.g., treatment vs. control) can be predicted from molecular measurements alone. 
This is achieved by training a machine-learning model specific to each cell type, to predict the experimental condition from which each individual cell originated.
The accuracy of each cell type-specific classifier is evaluated in cross-validation, providing a quantitative basis for cell type prioritization.

## System requirements

Augur relies on functions from the following R packages:

```
	dplyr (>= 0.8.0),
	purrr (>= 0.3.2),
	tibble (>= 2.1.3),
	magrittr (>= 1.5),
	tester (>= 0.1.7),
	Matrix (>= 1.2-14),
	sparseMatrixStats (>= 0.1.0),
	parsnip (>= 0.0.2),
	recipes (>= 0.1.4),
	rsample (>= 0.0.4),
	yardstick (>= 0.0.3),
	pbmcapply (>= 1.5.0),
	lmtest (>= 0.9-37),
	rlang (>= 0.4.0),
	glmnet (>= 2.0),
	randomForest (>= 4.6-14)
```

In addition, the [Seurat](https://satijalab.org/seurat/) and [monocle3](https://cole-trapnell-lab.github.io/monocle3/) packages must be installed for Augur to take Seurat and monocle objects as input, respectively.

Augur has been tested with R version 3.5.0 and higher.

## Installation

To install Augur, first install the devtools package, if it is not already installed: 

```r
> install.packages("devtools") 
```

Then, install the sparseMatrixStats package from GitHub using devtools:

```r
> devtools::install_github("const-ae/sparseMatrixStats")
```

Finally, install Augur from GitHub: 

```r
> devtools::install_github("neurorestore/Augur")
```

This should take no more than a few minutes.

## Usage

The main function of Augur, `calculate_auc`, takes as input a preprocessed features-by-cells (e.g., genes-by-cells for scRNA-seq) matrix, and a data frame containing metadata associated with each cell, minimally including the cell type annotations and sample labels to be predicted.
This means that in order to use Augur, you should have pre-processed your data (e.g., by read alignment and cell type assignment for scRNA-seq) across all experimental conditions. 
If batch effects are present in the data, these should be accounted for, e.g., using [Seurat](https://www.sciencedirect.com/science/article/pii/S0092867419305598) or [Harmony](https://www.nature.com/articles/s41592-019-0619-0), to avoid biasing cell type prioritization by technical differences or batch effects. 

To run Augur with default parameters on a genes-by-cells scRNA-seq matrix `expr`, and an accompanying data frame `meta`, with `cell_type` and `label` columns containing cell types and experimental conditions, respectively, use the `calculate_auc` function:

```r
> augur = calculate_auc(expr, meta)
```

If your columns have different names, you can specify these using the `cell_type_col` and `label_col` arguments:

```r
> augur = calculate_auc(expr, meta, cell_type_col = "cell.type", label_col = "condition")
```

Cell type prioritizations are stored in the `AUC` data frame - for example:

```r
> head(augur$AUC, 5)

# A tibble: 6 x 5
  cell_type                 auc
  <chr>                   <dbl>
1 cell type 1             0.752
2 cell type 2             0.729
3 cell type 3             0.674
  ...                     ...
```

Augur can also run directly on a Seurat object. For a Seurat object `sc`, with the `sc@meta.data` data frame containing `cell_type` and `label` columns, simply do:

```r
> augur = calculate_auc(sc)
```

The same code can be used if `sc` is a monocle3 object instead.

## Demonstration

To see Augur in action, load the simulated single-cell RNA-seq dataset that is bundled with the Augur package:

```r
> data("sc_sim")
```

This dataset consists of 600 cells, distributed evenly between three populations (cell types A, B, and C). Each of these cell types has approximately half of its cells in one of two conditions, treatment and control. The cell types also have different numbers of genes differentially expressed in response to the treatment. Cell type A has approximately 5% of genes DE in response to the treatment, while cell type B has 25% of its genes DE and cell type C has 50% of genes DE. 

The `sc_sim` object is a Seurat object with columns named `cell_type` and `label` in the `meta.data` slot, meaning we can provide it directly as input to Augur: 

```r
> head(sc_sim@meta.data)

      label cell_type
1   control CellTypeA
2 treatment CellTypeA
3 treatment CellTypeA
4   control CellTypeA
5   control CellTypeA
6 treatment CellTypeA
```

We run `calculate_auc`, and inspect the cell type prioritizations in the `AUC` item:

```r
> library(Augur)
> augur = calculate_auc(sc_sim)
> augur$AUC

# A tibble: 3 x 2
  cell_type   auc
  <chr>     <dbl>
1 CellTypeC 0.866
2 CellTypeB 0.763
3 CellTypeA 0.594
```

Augur has correctly recovered the simulated perturbation intensities. 

Running this example on a MacBook takes between 3 and 4 minutes. 
However, analyzing >20 real single-cell RNA-seq dataset, we found Augur takes a median of ~45 minutes.
In general, runtime scales close to linearly with the number of cell _types_.
By default, Augur runs on four cores, with each cell type analyzed on a different core.
To change the number of cores, use the `n_threads` argument.
For example, running Augur on eight threads: 

```r
> augur = calculate_auc(sc_sim, n_threads = 8)
```

... will run about twice as fast. 
